# A survey of spiking activity reveals a functional hierarchy of mouse corticothalamic visual areas

**DOI:** 10.1101/805010

**Authors:** Joshua H. Siegle, Xiaoxuan Jia, Séverine Durand, Sam Gale, Corbett Bennett, Nile Graddis, Greggory Heller, Tamina K. Ramirez, Hannah Choi, Jennifer A. Luviano, Peter A. Groblewski, Ruweida Ahmed, Anton Arkhipov, Amy Bernard, Yazan N. Billeh, Dillan Brown, Michael A. Buice, Nicolas Cain, Shiella Caldejon, Linzy Casal, Andrew Cho, Maggie Chvilicek, Timothy C. Cox, Kael Dai, Daniel J. Denman, Saskia E. J. de Vries, Roald Dietzman, Luke Esposito, Colin Farrell, David Feng, John Galbraith, Marina Garrett, Emily C. Gelfand, Nicole Hancock, Julie A. Harris, Robert Howard, Brian Hu, Ross Hytnen, Ramakrishnan Iyer, Erika Jessett, Katelyn Johnson, India Kato, Justin Kiggins, Sophie Lambert, Jerome Lecoq, Peter Ledochowitsch, Jung Hoon Lee, Arielle Leon, Yang Li, Elizabeth Liang, Fuhui Long, Kyla Mace, Jose Melchior, Daniel Millman, Tyler Mollenkopf, Chelsea Nayan, Lydia Ng, Kiet Ngo, Thuyahn Nguyen, Philip R. Nicovich, Kat North, Gabriel Koch Ocker, Doug Ollerenshaw, Michael Oliver, Marius Pachitariu, Jed Perkins, Melissa Reding, David Reid, Miranda Robertson, Kara Ronellenfitch, Sam Seid, Cliff Slaughterbeck, Michelle Stoecklin, David Sullivan, Ben Sutton, Jackie Swapp, Carol Thompson, Kristen Turner, Wayne Wakeman, Jennifer D. Whitesell, Derric Williams, Ali Williford, Rob Young, Hongkui Zeng, Sarah Naylor, John W. Phillips, R. Clay Reid, Stefan Mihalas, Shawn R. Olsen, Christof Koch

## Abstract

The mammalian visual system, from retina to neocortex, has been extensively studied at both anatomical and functional levels. Anatomy indicates the cortico-thalamic system is hierarchical, but characterization of cellular-level functional interactions across multiple levels of this hierarchy is lacking, partially due to the challenge of simultaneously recording activity across numerous regions. Here, we describe a large, open dataset (part of the *Allen Brain Observatory*) that surveys spiking from units in six cortical and two thalamic regions responding to a battery of visual stimuli. Using spike cross-correlation analysis, we find that inter-area functional connectivity mirrors the anatomical hierarchy from the *Allen Mouse Brain Connectivity Atlas*. Classical functional measures of hierarchy, including visual response latency, receptive field size, phase-locking to a drifting grating stimulus, and autocorrelation timescale are all correlated with the anatomical hierarchy. Moreover, recordings during a visual task support the behavioral relevance of hierarchical processing. Overall, this dataset and the hierarchy we describe provide a foundation for understanding coding and dynamics in the mouse cortico-thalamic visual system.

## Introduction

Mammalian vision is the most widely studied sensory modality. Probing its cellular substrate has yielded insights into how the stream of photons impinging onto the retina leads to conscious perception and visuo-motor behaviors. Yet the vast majority of our knowledge of physiology at the cellular level derives from small-scale studies subject to substantial uncontrolled variation, uneven coverage of neurons, and selective usage of stimuli. The field’s ability to validate models of visual function has been hampered by the absence of large-scale, standardized, and open *in vivo* physiology datasets (Olshausen & Field 2004; Carandini et al 2005). To address this shortcoming, we previously developed a 2-photon optical physiological pipeline to systematically survey visual responses (de Vries et al., 2019). Calcium imaging facilitates the monitoring of activity in genetically defined cell populations over the course of many sessions. However, it lacks high temporal resolution and single-spike sensitivity, and doesn’t easily allow distributed, simultaneous recordings from cortical and deep subcortical structures. We therefore developed a complementary pipeline that leverages Neuropixels probes to measure spiking activity in six cortical visual areas as well as two visual thalamic nuclei, LGN and LP.

The concept of hierarchy has informed ideas about the architecture of the mammalian visual system for more than 50 years (Hubel and Wiesel, 1962), and has inspired powerful multi-layered computational networks (Fukushima, 1980; Krizhevsky et al., 2012; Riesenhuber and Poggio, 1999). This hierarchy has been investigated most extensively in the macaque, from the LGN and primary visual cortex (V1) into frontal eye fields and beyond (Bullier, 2001; Chaudhuri et al., 2015; Felleman and Van Essen, 1991; Murray et al., 2014; Rockland and Pandya, 1979; Schmolesky et al., 1998; Yamins and DiCarlo, 2016).

The existence of such a hierarchy in the mouse, with its far smaller brain and dense cortical graph (Gămănuţ et al., 2018), is less clear (D’Souza and Burkhalter, 2017; Glickfeld and Olsen, 2017; Wang et al., 2012; Wang and Burkhalter, 2007). Yet given the utility of the laboratory mouse as a model organism, understanding the presence and extent of a hierarchy is of crucial importance. Harris, Mihalas et al applied multi-graph connectivity analysis to the *Allen Mouse Brain Connectivity Atlas* and inferred a shallow hierarchy in the full cortico-thalamic network, based on more than 1000 viral tracer injections aligned to a high-resolution 3D coordinate system (Harris et al., 2019). This analysis revealed a hierarchical ordering of visual areas in the mouse, with LGN at the bottom, and cortical area AM at the top. We sought to investigate whether this anatomical hierarchy is reflected in the spiking activity and functional properties of these visual areas.

Here, we describe a large-scale and systematic electrophysiological survey of spiking activity across visual cortico-thalamic structures in awake, head-fixed mice viewing diverse artificial and natural stimuli. We used Neuropixels silicon probes (Jun et al., 2017) to simultaneously record the electrical activity of hundreds of neurons with high spatial and temporal resolution (Allen et al., 2019; Steinmetz et al., 2018; Stringer et al., 2019). This dataset complements our previously released survey using optical recordings of calcium-evoked fluorescent activity in 60,000 cortical neurons (de Vries et al., 2019). Both datasets are part of the *Allen Brain Observatory*, a pipeline of animal husbandry, surgical procedures, equipment, and standard operating procedures (SOPs), coupled to strict, activity-and operator-independent quality control (QC) measures. All physiological data passing QC is made freely and publicly available via brain-map.org and the AllenSDK. Our initial characterizations of these pipelines treat each dataset independently; a detailed comparison of the results obtained via calcium imaging and electrophysiology is forthcoming.

Here, we have studied to what extent the flow of spikes follows the anatomical hierarchy by mining a *functional* dataset and relating it to a *structural* dataset. We first perform cross-correlation analysis between pairs of neurons to determine the relative timing of spiking activity across areas. We then demonstrate that a variety of functional metrics previously used to identify hierarchical processing support the cortico-thalamic hierarchy found neuroanatomically. Finally, recordings during active behavior suggest that one role of the hierarchy may be to amplify responses to behaviorally relevant stimulus changes (Brincat et al., 2018; Issa et al., 2018; Vinken et al., 2017).

### A survey of visually evoked spiking activity

Each mouse in the study proceeds through an identical series of steps, carried out by highly trained staff according to a set of SOPs (Figs. 1A and S1; see http://help.brain-map.org/display/observatory/Documentation). On the day of the experiment, we use cortical area maps derived from intrinsic signal imaging (ISI) to simultaneously target up to six Neuropixels probes to V1 and five higher-order visual cortical areas (LM, AL, RL, PM, AM) (Fig. 1B–D). As probes are inserted up to 3.5 mm into the brain, we regularly obtained concurrent recordings from two thalamic regions: the lateral geniculate nucleus (LGN) and the lateral posterior nucleus (LP, making up the visual pulvinar) (Fig. 1E), in addition to hippocampus and other areas traversed by the silicon probes. This configuration allowed us to sample the mouse visual system with unprecedented coverage, creating cellular-resolution activity maps across up to 8 corticothalamic visual areas at once, while also obtaining physiological measurements from nearby regions, such as hippocampus (Fig. 1F). The bulk of these recordings were made in C57BL/6J wildtype mice (*N* = 30), supplemented by recordings in three transgenic lines (*N* = 8 Pvalb-IRES-Cre x Ai32, *N* = 12 Sst-IRES-Cre x Ai32, and *N* = 8 Vip-IRES-Cre x Ai32), to facilitate the identification of genetically-defined inhibitory cell types via opto-tagging (Lima et al., 2009).

**Figure 1.**
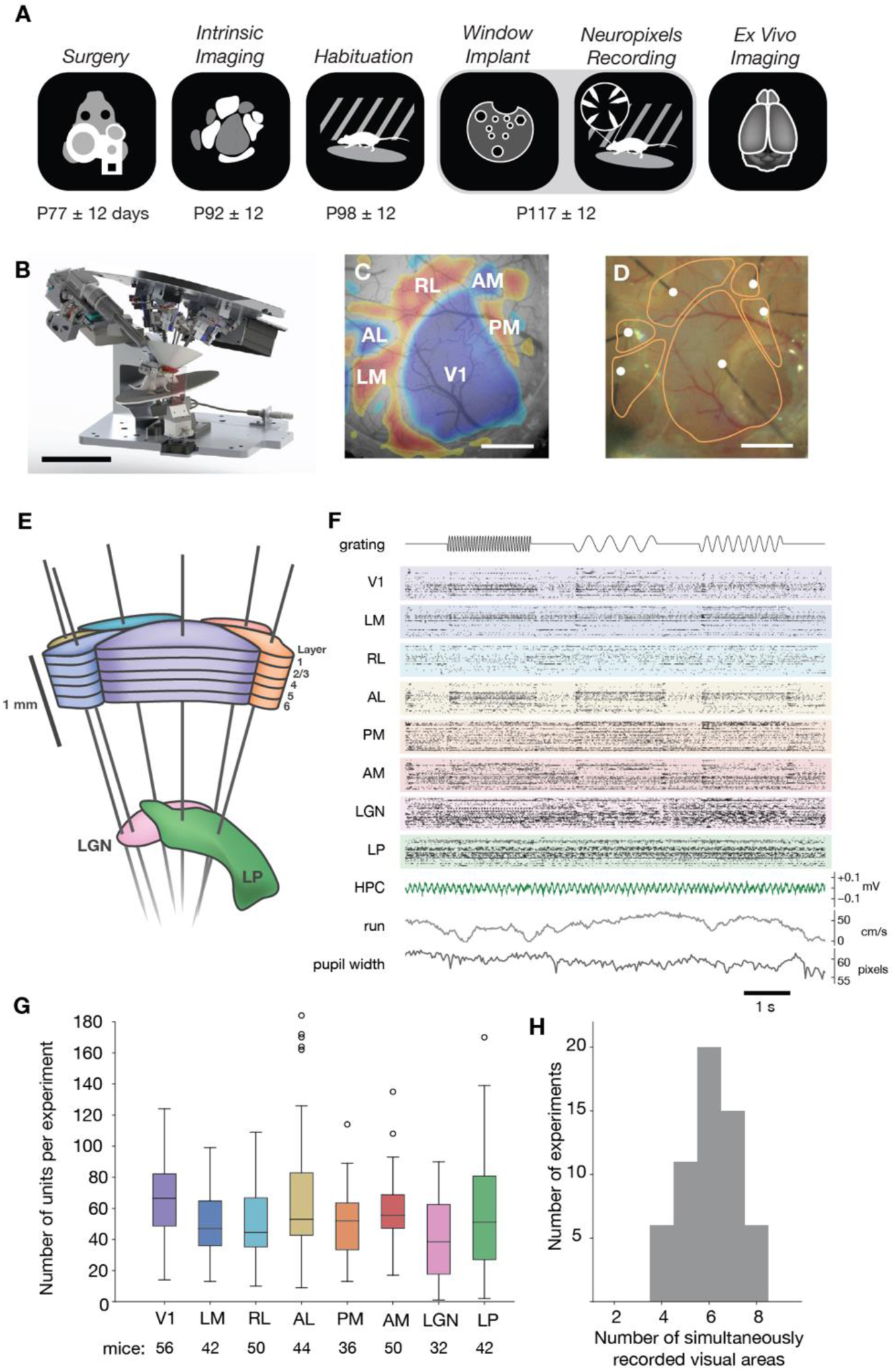
A standardized pipeline for extracellular electrophysiology in the mouse corticothalamic visual system. (**A**) Icons representing six major steps in the data collection pipeline, with the average age of mice at each step indicated below. (**B**) Rig for parallel recording from six Neuropixels probes. Scale bar = 10 cm. (**C**) Example retinotopic map used for targeting probes to six cortical visual areas. Scale bar = 1 mm. (**D**) Image of Neuropixels probes during an experiment, with area boundaries from (C) overlaid in orange. Probe tips are marked with white dots. Scale bar = 1 mm. (**E**) Schematic of target probe insertion trajectories through cortex into two thalamic visual areas, LGN and LP. (**F**) Example raster plot of 405 simultaneously recorded units from 8 visual areas. The phase of the drifting grating visual stimulus (15 Hz, 2 Hz, or 4 Hz), hippocampal local field potential, mouse running speed, and pupil width are also shown. (**G**) Box plot of the number of units recorded per area per experiment, after filtering based on ISI violations (<0.5), amplitude cutoff (<0.1), and presence ratio (>0.95) (see Methods and Figure S4 for quality metric definitions and distributions). (**H**) Histogram of the number of simultaneously recorded corticothalamic visual areas per experiment.

We implemented QC procedures to ensure consistent data across experiments (Fig. S2 and Methods), reducing the number of completed and analyzed experiments from 87 to 58. Extracellularly recorded units were identified and sorted via the automated Kilosort2 algorithm (Pachitariu et al., 2016; Stringer et al., 2019) followed by QC to remove units with artifactual waveforms, yielding a total of 99,180 units across experiments (Fig. S3). A variety of quality metrics were calculated to assess unit contamination and completeness, which are used to select units for further analysis (Fig. S4). Units were mapped to structures in the Common Coordinate Framework Version 3 (CCFv3) by imaging fluorescently labeled probe tracks with optical projection tomography (Fig. S5). After filtering units based on quality metrics, we simultaneously recorded from a mean of 682 ± 144 units per experiment, 128 ± 51 units per probe, and 56 ± 30 units per cortico-thalamic visual area (Fig. 1G). We sampled from a mean of 6.1 ± 1.1 cortico-thalamic visual areas in each experiment, with a subset of experiments including units from 8 visual areas simultaneously (Fig. 1H).

During each recording session, mice passively viewed a battery of natural and artificial stimuli (Fig. 2A and S6). In this study, we focused our analysis on a subset of these: drifting gratings (Fig. 2B), full-field flashes (Fig. 2C), and local Gabor patches (which are used to map spatial receptive fields) (Fig. 2D). Overall, units recorded in all 8 cortico-thalamic visual areas were highly visually responsive, with 60% displaying significant receptive fields (Fig. 2E and Fig. S7, categorical χ² test, *P* < 0.01). As a control, we searched for significant receptive fields in simultaneously recorded hippocampal regions (CA1, CA3, and dentate gyrus), and only found them in 1.4% of units. Mapping recorded units with a significant receptive field to their location within the CCFv3 and aggregating over experiments, we recapitulated the previously described retinotopic map organization in each area (Fig. 2F) (Bennett et al., 2019; Garrett et al., 2014; Román Rosón et al., 2019), supporting the accuracy of spatial registration in the CCFv3.

**Figure 2.**
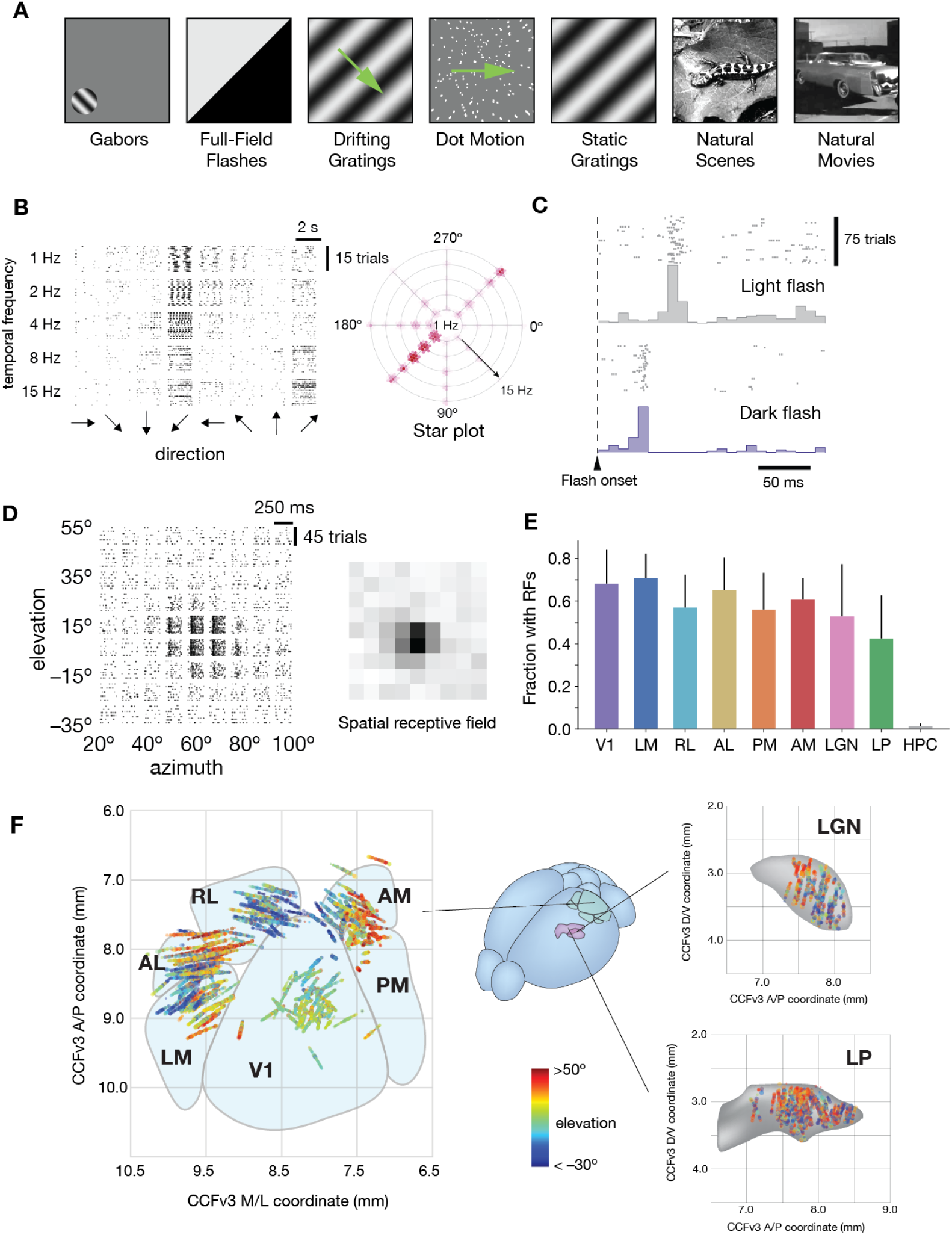
High-throughput mapping of visual response properties. (**A**) During Neuropixels recordings, mice are exposed to up to seven types of natural and artificial stimuli. (**B**) Raster plots of spike times for 40 unique conditions of a drifting grating stimulus, for an example V1 unit. The single-trial responses are used to construct a “star plot,” which efficiently summarizes the unit’s tuning properties. (**C**) Raster plot of spike times for two conditions of the full-field flash stimulus, for the same unit in B. A peri-stimulus time histogram summarizes the response across trials. (**D**) Raster plot of spike times for 81 conditions of the Gabor stimulus for the same unit as in B and C. Summing the spike counts across trials at each location produces a spatial receptive field, shown on the right. (**E**) Mean fraction of units with significant receptive fields (RFs) across 8 visual areas, with hippocampus included as a control. Error bars represent standard deviation across experiments. (**F**) Each unit with a significant receptive field is represented by a dot at its spatial location in the mouse Common Coordinate Framework. Color represents the elevation of the receptive field center, revealing smoothly varying maps of visual space when aggregating across experiments. Area boundaries are approximate.

### A functional hierarchy of visual areas

The work of (Harris, Mihalas, et al. 2019) assigned a hierarchy score to each of the visually responsive areas from which we recorded. This score is derived using an optimization algorithm that considers the set of distinct axonal termination patterns of connectivity between areas— deeming each as either feedforward versus feedback connections—and finds the most self-consistent network architecture (Fig. 3A; Supplementary Methods). The LGN sits at the bottom of the hierarchy, followed by its major target structure, V1. Four higher-order visual areas (LM, RL, LP, and AL) reside at intermediate levels, with areas PM and AM occupying the top level of the areas we studied here (Fig. 3A,D).

**Figure 3.**
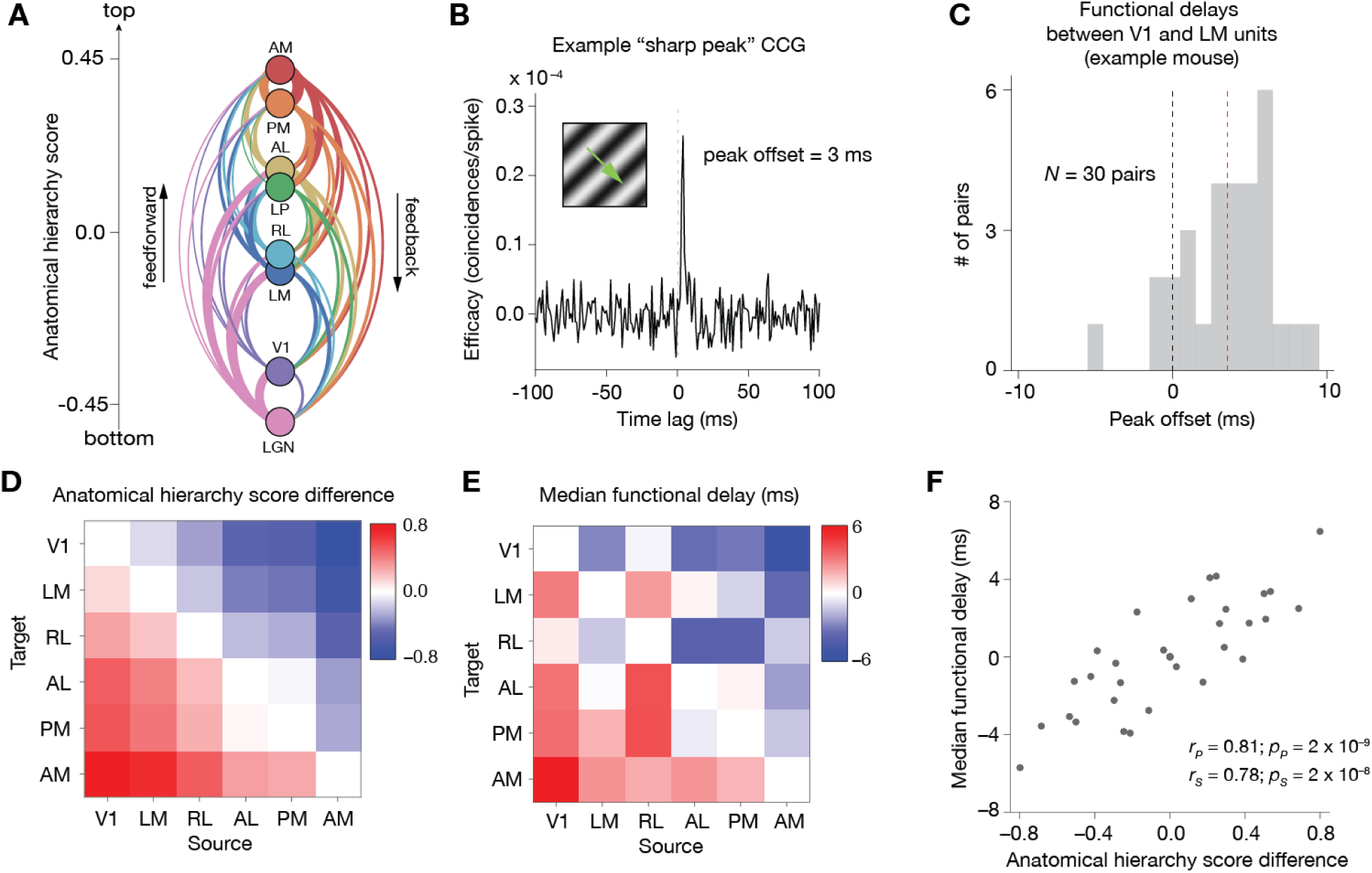
Functional connectivity recapitulates the anatomical hierarchy. (**A**) Relative anatomical hierarchy scores from Harris, Mihalas et al. (2019). Each area’s hierarchy score is based on the ratio of feed-forward vs. feedback projection patterns (colored lines) between itself and the other 7 areas. (**B**) Method for measuring cross-area spiking interactions between pairs of units. “Sharp peaks” in the jitter-corrected CCG are those with a peak amplitude >7x the standard deviation of the flanks. The peak offset, or functional delay, is defined as the difference between the CCG peak time and the CCG center (at 0 ms). (**C**) Distribution of functional delay between V1 and LM reflected by pairwise CCG peak offset in one example mouse (*N* = 30 pairs; median = 3.9ms). (**D**) Re-plotting of anatomical hierarchy scores from (A), showing the difference in score between all pairs of cortical areas. Statistical testing (Wilcoxon Rank-sum test) revealed that all areas have significant different hierarchical score, except for RL and LM. (**E**) Combined median of functional delay across mice (*N* = 25 mice in total) for each pair of cortical areas. Statistical testing (Wilcoxon Rank-sum test) revealed that the peak offset distribution of neighboring-areas were significantly different from within-area, except for AL-PM. (**F**) Correlation between the median functional delay and the difference in hierarchy scores, indicating a link between structure and function (Pearson’s *r* = 0.81, *P* = 2e-9).

To compare this anatomical hierarchy to a possible functional hierarchy measured in the spike recordings, we computed a directional metric of functional connectivity, the spike cross-correlogram (CCG), between units (Jia et al., 2013; Smith and Kohn, 2008; Zandvakili and Kohn, 2015). This analysis focused on activity during periods of drifting grating presentation. For each pair of recorded units, we identified significant functional interactions as determined by a short latency (<10 ms) sharp peak in the jitter-corrected CCG (Fig. 3B), which removed slow timescale correlations larger than the jitter window (25 ms). This selection criterion yielded 56,874 pairs of units out of 12,908,146 total possible pairs (0.44%). These fast-timescale spiking interactions can provide a measure of the functional hierarchical relationship between areas, based on the measured relative spike timing between pairs of units (Fig. 3B example pair, peak offset = 3 ms). If units in one region tend to lead the spiking activity of higher regions, the distribution of peak offsets would deviate from 0. For example, the peak offset distribution of CCG pairs between V1 and LM showed a significant positive delay when compared to V1–V1 distribution (example mouse, Fig 3C; *N* = 30 pairs, *P* = 2.6e-8, Wilcoxon Rank-sum test), indicating V1 neurons spike earlier than LM and thus are lower in the hierarchy. As we had only limited experiments with measurable sharp CCG peaks between thalamic and cortical areas (LGN–cortex: 1 mouse; LP–cortex: 2 mice), we restricted our CCG-based hierarchy analysis to cortico-cortical interactions.

We quantified the distribution of sharp CCG peak time lags for all functionally connected pairs of units across each pair of cortical areas in each mouse and combined the median of peak offset distributions across mice (Fig. 3E; *N* = 25 mice; see Fig. S8 for complete peak offset distributions between all areas across all mice). V1 units consistently fired action potentials earlier than units in other areas (Fig. 3E, left column). In contrast, area AM consistently fired later than other regions, indicating this area resides at the uppermost levels of the hierarchy (Fig. 3E, right column). Quantifying the functional delay between all pair-wise sets of areas revealed an organization remarkably similar to the anatomical hierarchy (Fig. 3D). The correlation between the anatomical hierarchy score and the median temporal delay between all regions was high (Fig. 3F; Pearson’s *r* = 0.81, *P* = 2e–9; Spearman’s *r* = 0.78, *P* = 2e-8), indicating that unit spiking activity follows a functional hierarchy closely mirroring the anatomical structure of the visual cortico-thalamic system.

We next assessed how this ordering of areas correlated with four classical measures of functional hierarchy in primates (Bullier, 2001; Chaudhuri et al., 2015; Schmolesky et al., 1998). First, we quantified the temporal latency of responses to full-field flashes. Whereas cells in each area have broadly distributed onsets (Fig. 4A,B), consistent with primate results (Schmolesky et al., 1998), the mean visual latency of each area was correlated with its anatomical hierarchy score (Fig. 4C; Pearson’s *r* = 0.95, *P* = 0.00025). Statistical testing revealed significantly different latencies for all pairs of areas, except for LGN–V1, RL–LP, LP–AL, and AM–PM (Fig. S9A,B).

**Figure 4.**
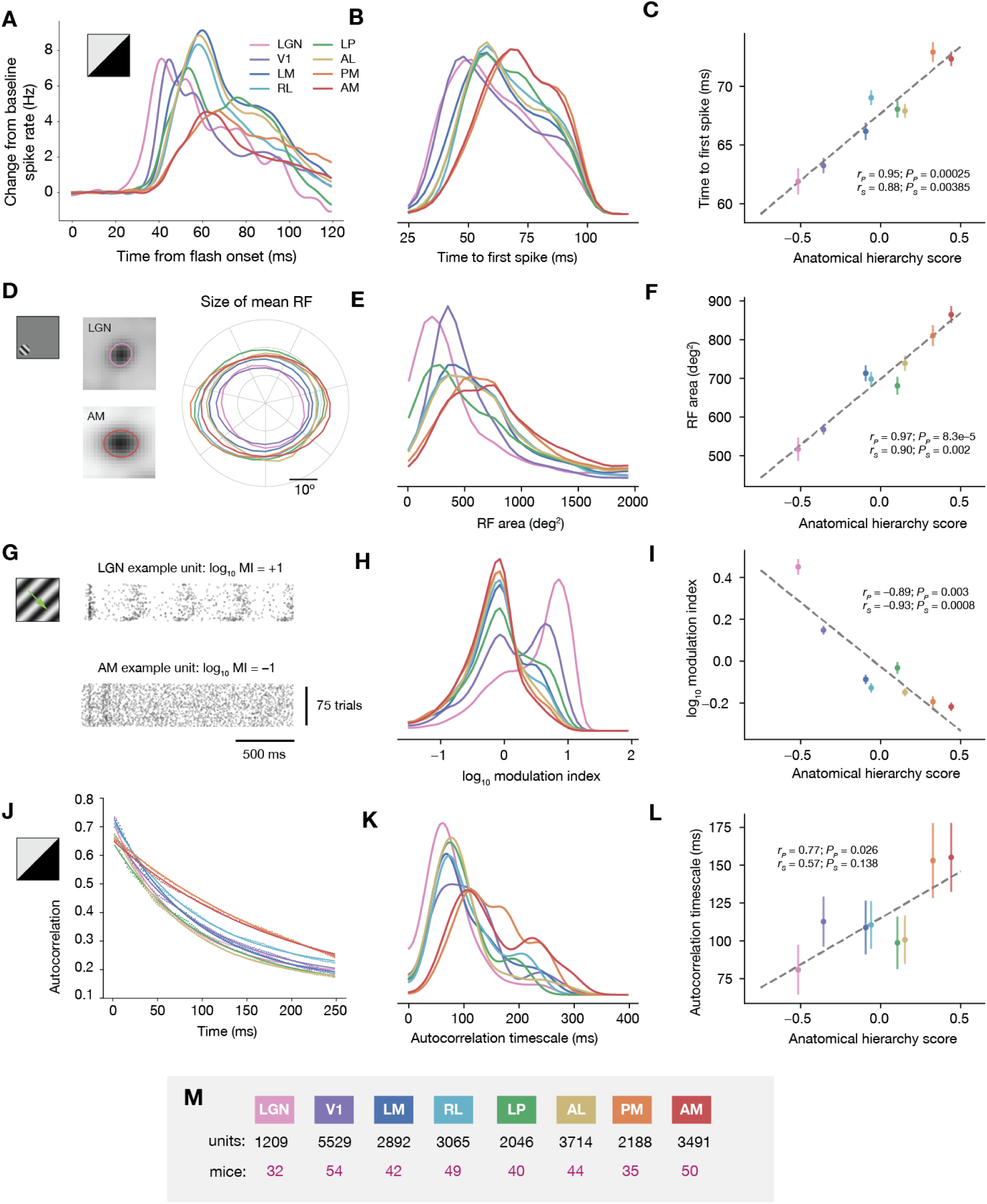
Four measures of hierarchical processing applied to the mouse visual system. (**A**) Mean peri-stimulus time histogram (PSTH) of the spiking response to a full-field flash stimulus across 2 thalamic and 6 cortical regions, with the baseline firing rate subtracted for each area. (**B**) Distribution of first spike times in response to the flash stimulus across all units in each of 8 areas. (**C**) Correlation between mean time to first spike and hierarchy score obtained from anatomical tracing studies. (**D**) Outlines of the extent of the mean receptive field (RF) for each area, at 50% of the peak firing rate. Exemplar mean receptive fields for LGN and AM are shown on the left. (**E**) Distribution of receptive field sizes across all units in each of 8 areas. (**F**) Correlation between mean receptive field size and anatomical hierarchy score. (**G**) Raster plots showing the response of exemplar units in LGN and AM to a 2 Hz drifting grating stimulus. Modulation index (MI) is higher in units that fire at the same temporal frequency as the grating. (**H**) Distribution of MI across all units in each of 8 areas. (**I**) Correlation between mean MI and anatomical hierarchy score. (**J**) Mean autocorrelation values for 8 areas in the 250 ms period following the onset of a full-field flash stimulus. An exponential fit is used to determine the autocorrelation timescale. (**K**) Distribution of autocorrelation timescales for all units in each of 8 areas. (**L**) Correlation between mean autocorrelation timescale and anatomical hierarchy score. (**M**) Figure legend, indicating color of each area, total number of units per area, and total number of mice per area. All error bars represent 95% bootstrap confidence intervals. See Fig S4B for unit selection criteria.

Second, the size of spatial receptive fields typically increases when ascending the visual processing stream (Freeman et al., 2013; Hubel, 1988; Lennie, 1998; Wang and Burkhalter, 2007), likely due to the pooling of convergent inputs from neurons in lower regions. We measured receptive fields using a localized Gabor stimulus (Figure 4D), and found a systematic increase in receptive field size with anatomical hierarchy score (Fig. 4D-F; Pearson’s *r* = 0.97, *P* = 8.3e-5). Statistical testing revealed significantly different receptive field sizes for all pairs of areas, except for LM–RL (Fig. S9C,D).

Third, Hubel and Wiesel (1962) described ‘simple’ and ‘complex’ cells in V1. Complex cells are thought to result from the integration of inputs from simple cells with different preferred spatial phases, and can therefore be identified by the lack of phase-dependent responses to a drifting grating stimulus (Hubel and Wiesel, 1965, 1962; Matteucci et al., 2019; Riesenhuber and Poggio, 1999). The fraction of cells with phase-dependent grating responses, which are common in the retina and decrease up the visual hierarchy, is a useful measure of hierarchical level. We quantified this with a modulation index (MI) that robustly reflects phase-dependent responses to drifting gratings (Matteucci et al., 2019; Wypych et al., 2012). MI measures the difference in power of the visually evoked response at a unit’s preferred stimulus frequency versus the average power spectrum. MI > 3 corresponds to strong modulation of spiking at the stimulus frequency (indicative of simple-cell-like responses), whereas smaller MI values indicate less modulation by stimulus temporal frequency (indicative of complex-cell-like responses) (Matteucci et al., 2019). Compatible with more simple-like processing, MI was higher in LGN and V1, whereas the higher order areas showed considerably less phase-dependent modulation (Figure 4G-I, Pearson’s *r* = –0.89, *P* = 0.003). Statistical testing revealed significantly different modulation indices for all pairs of areas, except for RL–AL and AM–PM (Fig. S9E,F).

Finally, previous work in the primate brain demonstrated that the ‘timescale’ of neural activity increases at higher levels of the cortical hierarchy (Chen et al., 2015; Murray et al., 2014). We quantified the temporal scale for each area in our study by fitting an exponential decay to its mean spike-count autocorrelation function following a full-field flash stimulus (Fig. 4J). We found that higher-order areas had a longer timescale and thus integrate over longer temporal windows than lower stages, an important signature of multi-layer processing (Fig. 4J-L; Pearson’s *r* = 0.77, *P* = 0.026). Statistical testing revealed significantly distinct timescales for LGN–V1, LGN–RL, and for AM and PM vs. all other areas (Fig. S9G,H).

Together, these four response metrics and the functional connectivity analysis support the existence of a functional hierarchy spanning the mouse corticothalamic visual system. These metrics are not dependent on overall firing rate, which does not correlate with hierarchy score (Fig. S9I-J).

The role of this hierarchical structure should ultimately be related to the behavioral and cognitive operations implemented by the system. Higher levels of the hierarchy are positioned to increasingly integrate sensory input with behavioral goals. To test whether the hierarchy we found correlates with behaviorally relevant processing, we performed additional experiments beyond our passive viewing survey to measure spiking activity across the visual hierarchy while mice actively performed a visual change detection task (4527 units from 14 mice) (Garrett et al., 2019). In this go/no-go task, mice report when a visual stimulus (here, natural scenes; Fig. 5A) changes identity by licking a reward spout.

**Figure 5.**
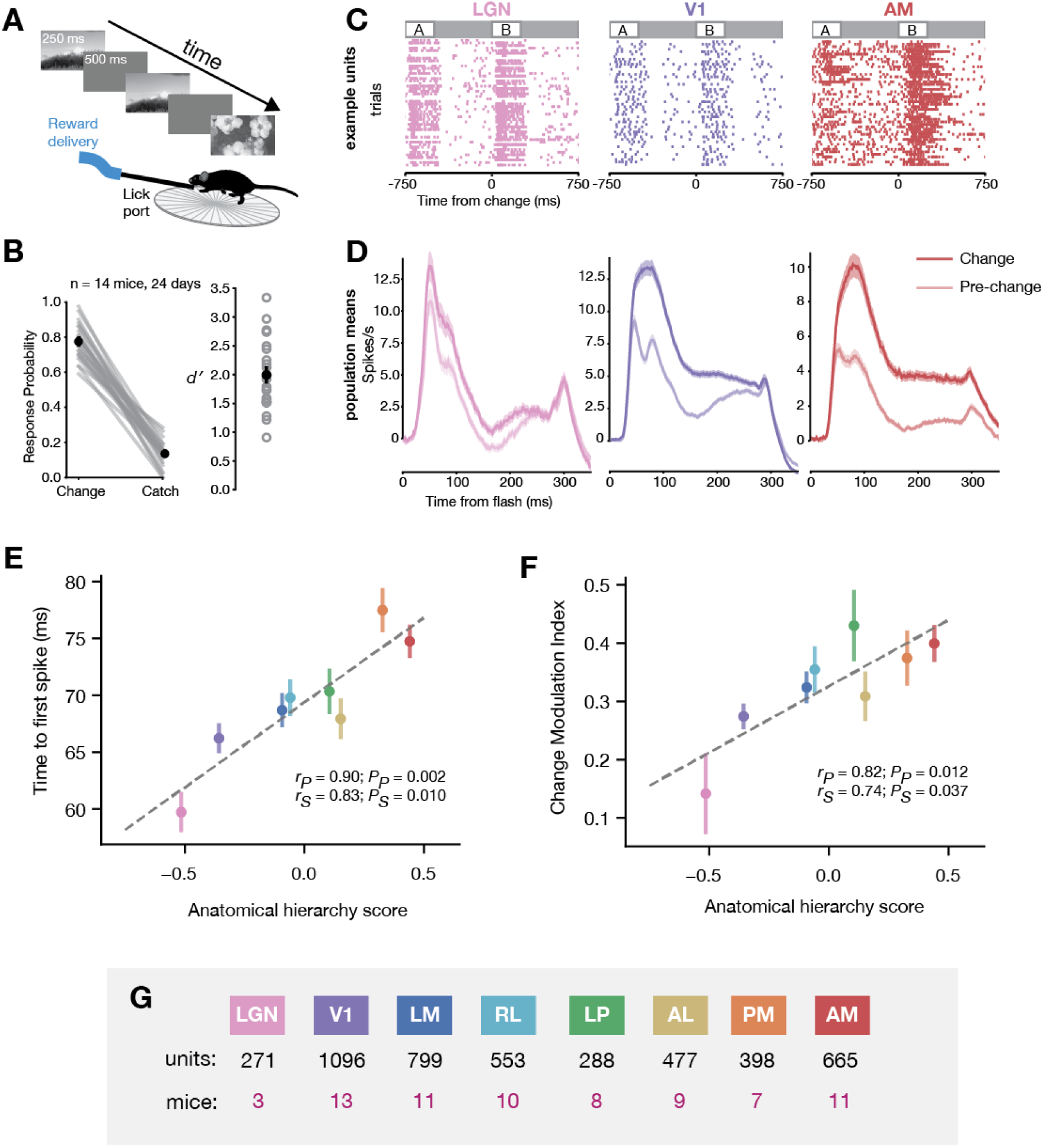
Higher-order areas signal behaviorally relevant changes in image identity more strongly than lower-order areas. (**A**) Mice were rewarded when they correctly detected change in the identity of a natural scene shown for 250 ms, separated by 500 ms blank screen. (**B**) After training, 10 mice had high hit rates and low false alarm rates, with an average *d′* of 1.9 +/-0.2. (**C**) Rasters showing spiking of exemplar units from LGN, V1, and AM during 50 trials of the change detection task. (**D**) Population peri-stimulus time histograms averaged over all units in LGN, V1, and AM. For each area, the response to the change image is shown as a darker line, while the response to the pre-change image is shown as a lighter line. The change modulation index is defined as the normalized difference between the firing rate during the pre-change image and the change image. (**E**) Correlation between mean time to first spike and anatomical hierarchy score across all 8 areas during the change detection task. (**F**) Correlation between mean change modulation index and anatomical hierarchy score across all 8 areas. Responses to the stimulus change are always greater than the pre-change flash (CMI > 0) and this difference increases from LGN to AM. (**G**) Figure legend, indicating color of each area, total number of units per area, and total number of mice per area. Error bars in E and F represent 95% bootstrap confidence intervals.

During the Neuropixels recordings, mice performed the task with high hit rates and low false alarm rates (mean hit rate = 0.70, mean false alarm rate = 0.12, mean *d′ =* 1.9 ± 0.2; Fig. 5B). Units recorded during the task had clear visually evoked spiking responses to the flashed visual stimuli and showed greater evoked spike rates when the visual stimulus changed identity (from A to B at *t* = 0 in Fig. 5C-D). As in the passive viewing mice, latency to first spike in response to the stimulus was correlated with hierarchy score (Fig. 5E). These latencies during active behavior were highly correlated with latencies measured in the passive viewing condition (Pearson’s *r* = 0.95, *P* = 0.003). Since this task requires mice to detect stimulus changes, for each unit we computed a ‘change modulation index’ to capture the differential response to repeated versus changed images (this metric varies from −1 to 1 with 0 representing no modulation). The mean change modulation was positive for each area, indicating that a change in image identity elicits stronger responses compared to the same image presentation. Importantly, change modulation systematically increased along the hierarchy from LGN to AM (Fig. 5F; Pearson’s *r* = 0.82, *P* = 0.013). Other aspects of neural activity during the task, such as the pre-change spike rate, the change response spike rate, and the baseline firing rate were not correlated with hierarchy score (Fig. S10E-G). Statistical testing revealed that all pairs of areas have significantly different change modulation indices, except for LM–AL, RL–PM, RL–AM, and PM–AM (Fig. S10D). This suggests that change-related signals are amplified at higher levels of the visual hierarchy.

## Discussion

One long-term goal of the Allen Institute is to systematically survey neuronal activity in the visual corticothalamic complex, responding to a battery of commonly used visual stimuli in awake mice in a way that is minimally biased, maximally reproducible, and freely accessible to all (Koch and Reid, 2012). We previously presented one such survey, based on two-photon calcium imaging, that captured cellular activity in six cortical regions in various transgenic animals (de Vries et al., 2019). We here complement this *Allen Brain Observatory* database at www.brain-map.org with a survey of spiking activity measured using high-density silicon Neuropixels probes (Jun et al., 2017). We recorded from the same cortical regions as in the two-photon imaging survey, in addition to thalamic visual areas LGN and LP. (We also recorded units from hippocampus and nearby areas due to the 3.5 mm span of our Neuropixels recordings; these also are included in our open data release.) In agreement with the foundational studies of mouse visual cortex (Andermann et al., 2011; Marshel et al., 2011; Niell and Stryker, 2008), we find a plethora of units across the 8 regions with robust visual responses, with 60% of units displaying significant visual receptive fields (Fig. 2E).

For this first report on our electrophysiological survey, we focused on one important aspect of this rich dataset – exploiting the dynamic flow of spikes between brain areas to infer functional hierarchical processing in the visual cortico-thalamic system and relating this to quantitative measures of its anatomical hierarchy. Based on anterograde viral tracing with Cre-dependent AAV in 1,256 experiments in 50 distinct mouse lines, Harris et al (2019) derived anatomical rules describing cortico-cortical, cortico-thalamic, and thalamo-cortical projections into and out of 37 cortical and 24 thalamic regions via their layer-specific axonal termination patterns. Using an optimization approach that labels connections as either feedforward or feedback to find the most self-consistent network, the algorithm assigns a hierarchy score to every region (Fig. 3A). The study clearly demonstrates that the full corticothalamic system of the mouse is hierarchically organized, but the difference between the lowest and highest rungs is only a few full levels, due to parallel and short-cut projections among areas; thus, this system is organized as a ‘shallow’ hierarchy.

From the complete corticothalamic anatomical hierarchy of Harris et al (see Fig. 6d in their paper), we extracted the six cortical and two thalamic visual areas we targeted with Neuropixels probes. Our analysis indicates these areas represent at least seven distinct levels starting with LGN and V1, followed by LM/RL, LP, AL, and finally PM and AM at the highest rungs (Fig. 3A), consistent with previous anatomical hierarchy schemes in rodents (Coogan and Burkhalter, 1993; D’Souza et al., 2016). It follows that neurons at a lower level of this hierarchy should spike earlier than neurons at higher levels (this will only be true on average as there are many feedback pathways, neurons with a diversity of time constants, and other sources of heterogeneity). Accordingly, for those pairs of units in any two regions with overlapping receptive fields, we compute the CCG to identify short latency peaks and extract the relative spike timing between pairs of neurons (Fig. 3E; Fig. S8); note that the range of temporal lags involved, ±10 ms, may include both mono-and multi-synaptic connections. This uncovers a striking correspondence between the anatomical and functional network organization—the bigger the difference in the anatomical hierarchy score of two areas, the larger the median time lag of spikes between these regions (Fig. 3F).

We quantified visual responses in the ascending areas of this visual hierarchy by computing four previously used measures of hierarchical processing – response latency, receptive field size, degree of phase modulation by a drifting grating stimulus, and autocorrelation timescale. All four measures either increased or decreased systematically across these eight visual regions, and pairwise statistical tests suggest that each of these measures can independently differentiate between distinct hierarchical levels (Fig. 4; Fig. S9).

Even though these functional metrics are correlated with the anatomical hierarchy score, it is important to note that describing the system with a single variable is a simplification, meant only to reflect a first-order characterization of its organization. While we have simultaneously recorded spiking activity from more mouse visual areas than any previous study, we only sampled six of the 16 mouse cortical visual areas (Zhuang et al., 2017). Moreover, there is substantial overlap in the distribution of response properties across areas—for example, many units in AM spike earlier than the slowest units in LGN (Fig. 4B). This suggests much processing occurs in parallel, in addition to the general hierarchical sequence in response to sensory drive. The primate visual system is organized into distinct streams (Maunsell, 1992; Ungerleider and Mishkin, 1982), and there is anatomical and functional evidence for parallel streams in mice (Smith et al., 2017; Wang et al., 2012). Future studies can uncover how distributed activity dynamics emerge from connectivity within these hierarchical and parallel circuits.

The functional hierarchy we establish provides a general framework for investigating how it is used to solve behaviorally relevant tasks. As a first step in this direction, we carried out experiments to assess whether this same hierarchy is visible in measures reflecting signal processing during behavior (Fig 5.). We found that the relative response to a rewarded changed image increased systematically from LGN to AM (Fig. 5F). This increase in change-related signals suggests that unexpected stimuli are amplified by successive levels of the hierarchy, consistent with evidence from an oddball paradigm in rat higher-order cortex (Vinken et al., 2017). These results are compatible with general theories of hierarchical predictive processing, which posit that unexpected signals are preferentially passed to higher processing stages (Dürschmid et al., 2016; Grimm et al., 2011; Issa et al., 2018; Keller and Mrsic-Flogel, 2018).

A major challenge in neuroscience is to understand how spiking activity flows through distributed brain networks to mediate cognition and behavior. The cortex is widely and densely connected (Gămănuţ et al., 2018; Oh et al., 2014), with diverse cell-type–specific anatomical pathways. The concept of hierarchy is one important first-order organizing principle for understanding form and function in the brain (Sporns, 2010). Reinforcing the anatomical and functional evidence for a mouse cortical hierarchy, cell type composition and gene expression systematically change across the global hierarchy in the mouse cortex (Fulcher et al., 2019; Kim et al., 2017; Tasic et al., 2018). Nonetheless, the cortex also displays additional levels of organization including functional sub-modules and parallel processing streams. These aspects must be incorporated to establish a more complete mapping between cortical structure and function (Han et al., 2018; Lennie, 1998; Ungerleider and Mishkin, 1982; Wang et al., 2012).

## Acknowledgements

We thank the Allen Institute founder, Paul G. Allen, for his vision, encouragement and support.

## Author contributions

Conceptualization: C.K., S.R.O, J.H.S., X.J., S.G., C.B., S.M., D.D., S.dV., M.B., C.R.

Supervision: C.K., S.R.O, J.H.S., P.A.G., C.R., C.F., S.M., H.Z., S.D.

Investigation, validation, methodology, and formal analyses: J.H.S., S.D., G.H, T.R., X.J., S.G., C.B., S.R.O., J.L., N.G., A.A., A.B., Y.B., M.B., L.C., N.C., S.C., A.C., T.C., S.dV., D.D., R.D., D.F., E.G., R.H., B.H., R.I., I.K., J.K., S.L., J.L., P.L., J.H.L, A.L., Y.L., F.L, K.M., L.N., T.N., R.N., G.O., M.O., J.P., M.R., D.R., M.R., S.S., C.L., M.S., D.S., J.S., D.W., A.W., R.A., D.B., M.C., E.L., K.R., K.N., B.S., E.J., K.J., J.M., K.N., M.G., D.O., J.H., J.D.W.,

Software: J.H.S., X.J., N.G., K.D., S.G., C.B., S.dV., M.P., D.O., J.K., N.C., H.C., D.R., D.W., J.G., M.B., P.L.

Data Curation: J.H.S., N.G., X.J., K.D., D.F., J.G., R.H., W.W., R.Y.

Project administration: C.T., S.N., L.C., L.E., N.H., J.W.P.

Visualization: J.H.S, X.J., S.G, C.B., H.C., S.D.

Original draft written by C.K., S.R.O., J.H.S., X.J. with input and editing from S.M., H.C., C.B., S.G.

All co-authors reviewed the manuscript.

---

**Supplementary Figure 1.**
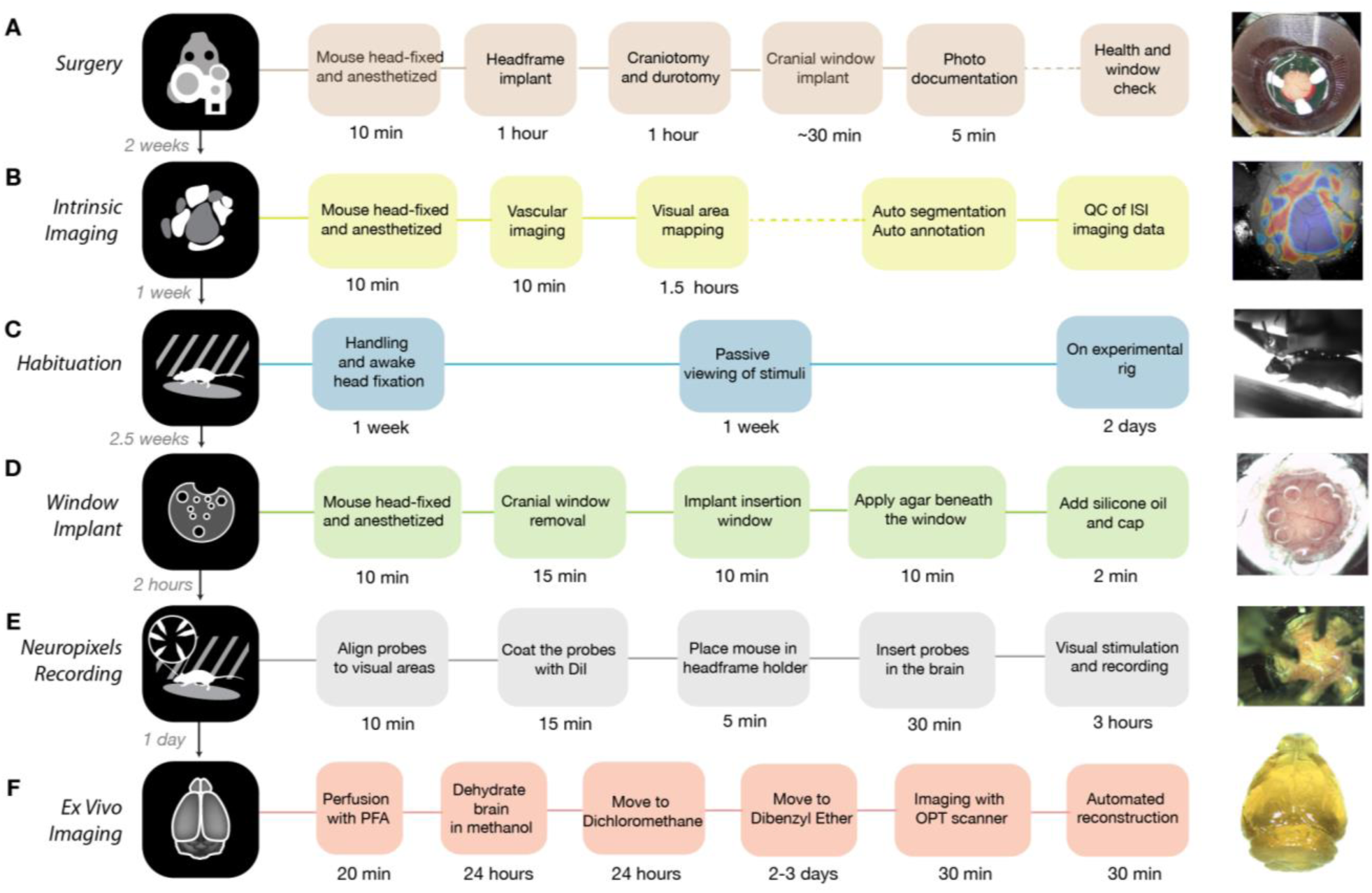
Pipeline procedures. Summary of procedures involved in each step of the pipeline.

**Supplementary Figure 2.**
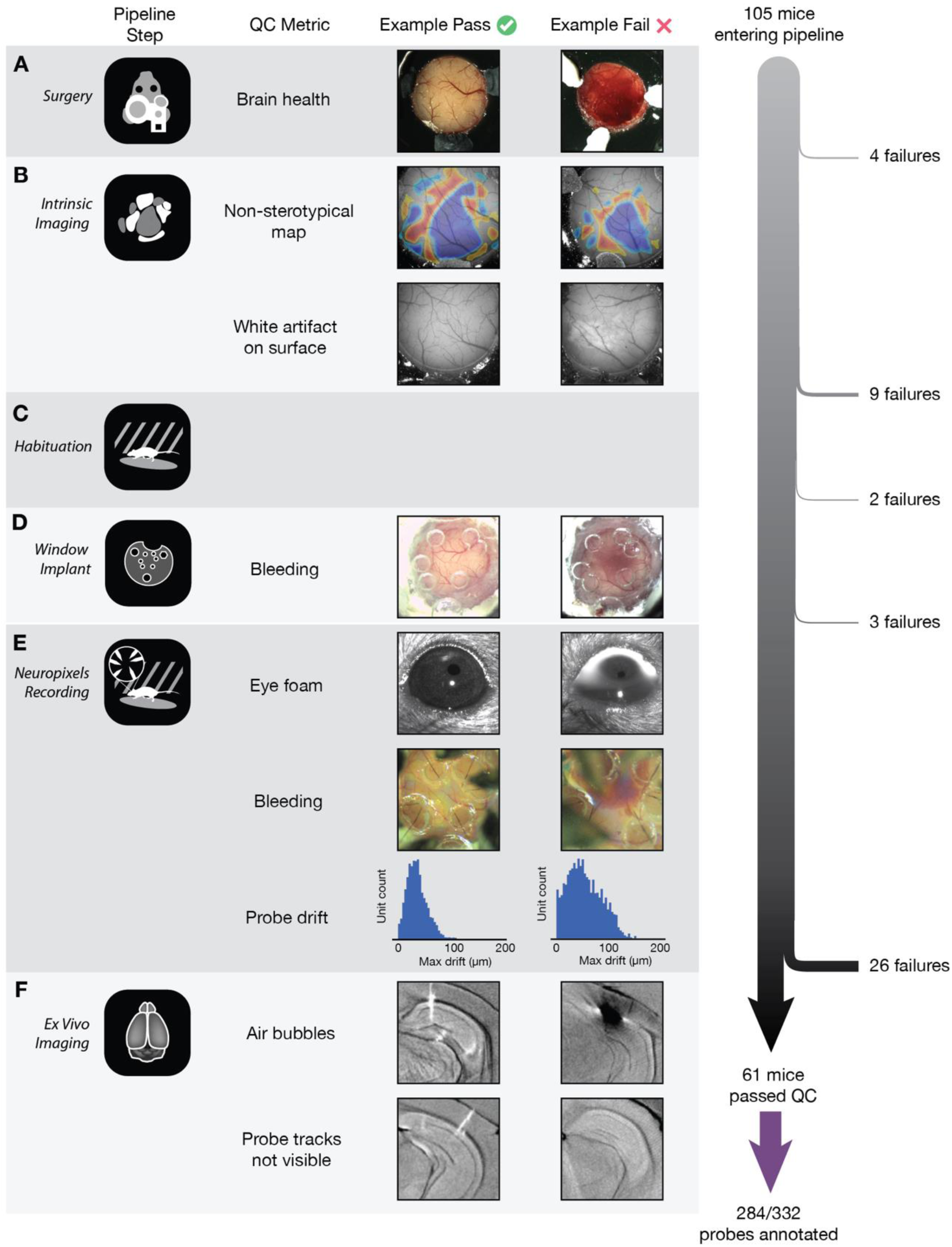
Pipeline quality control. Major QC metrics for each pipeline step, with examples of passing and failing experiments. The number of mice failing QC at each stage is shown on the right.

**Supplementary Figure 3.**
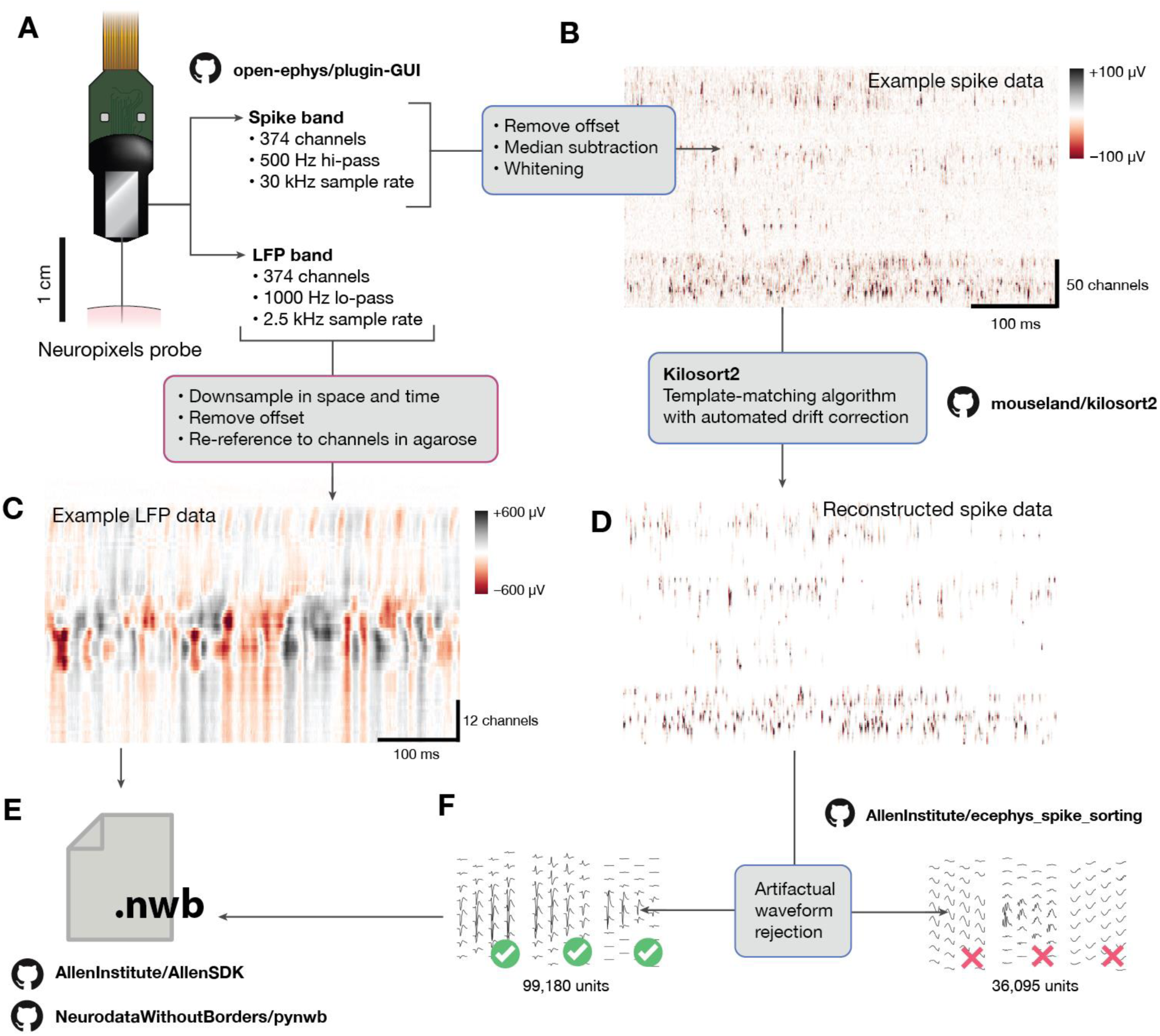
Data processing pipeline. (**A**) Data from the Neuropixels probe is split at the hardware level into two separate streams for each electrode: spike band and LFP band. (**B**) The spike band passes through offset subtraction, median subtraction, and whitening steps prior to sorting. The resulting data can be viewed as an image, with dimensions of time and channels, and colors corresponding to voltage levels. (**C**) The LFP data is down-sampled to 1.25 kHz and 40 µm channel spacing prior to packaging. (**D**) We use the Kilosort2 to match spike templates to the raw data. The output of this algorithm can be used to reconstruct the original data using information about template shape, times, and amplitudes. (**E**) The spike and LFP data are packaged into Neurodata Without Borders (NWB) files. (**F**) The outputs of Kilosort2 are passed through a semi-automated QC procedure to remove units with artifactual waveforms. Only units with obvious spike-like characteristics are used for further analysis.

**Supplementary Figure 4.**
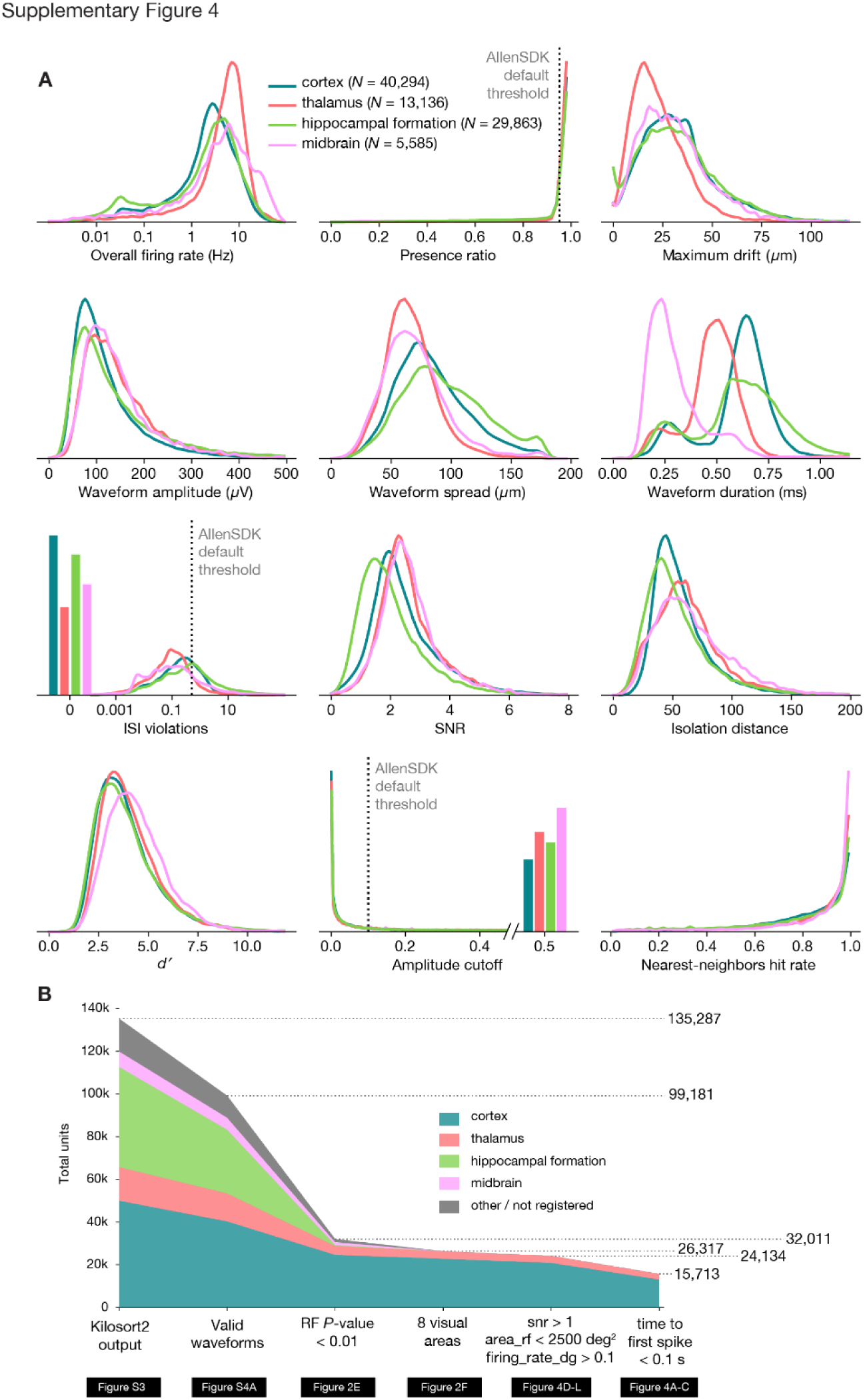
Unit quality metrics. **(A)** Density functions for 12 unit QC metrics, plotted for units in cortex, hippocampus, thalamus, and midbrain, aggregated across experiments. Default AllenSDK thresholds are shown as dotted lines. **(B)** Unit selection flowchart for generating manuscript figures. Note that we do not use the default AllenSDK filters in this work, but instead use an RF *P*-value of 0.01 as the primary metric for selecting units for analysis. CCFv3 structure labels used for region identification are as follows: cortex (VISp, VISl, VISrl, VISam, VISpm, VISal, VISmma, VISmmp, VISli, VIS), thalamus (LGd, LD, LP, VPM, TH, MGm, MGv, MGd, PO, LGv, VL, VPL, POL, Eth, PoT, PP, PIL, IntG, IGL, SGN, VPL, PF, RT), hippocampal formation (CA1, CA2, CA3, DG, SUB, POST, PRE, ProS, HPF), midbrain (MB, SCig, SCiw, SCsg, SCzo, SCop, PPT, APN, NOT, MRN, OP, LT, RPF), other / nonregistered (CP, ZI, grey).

**Supplementary Figure 5.**
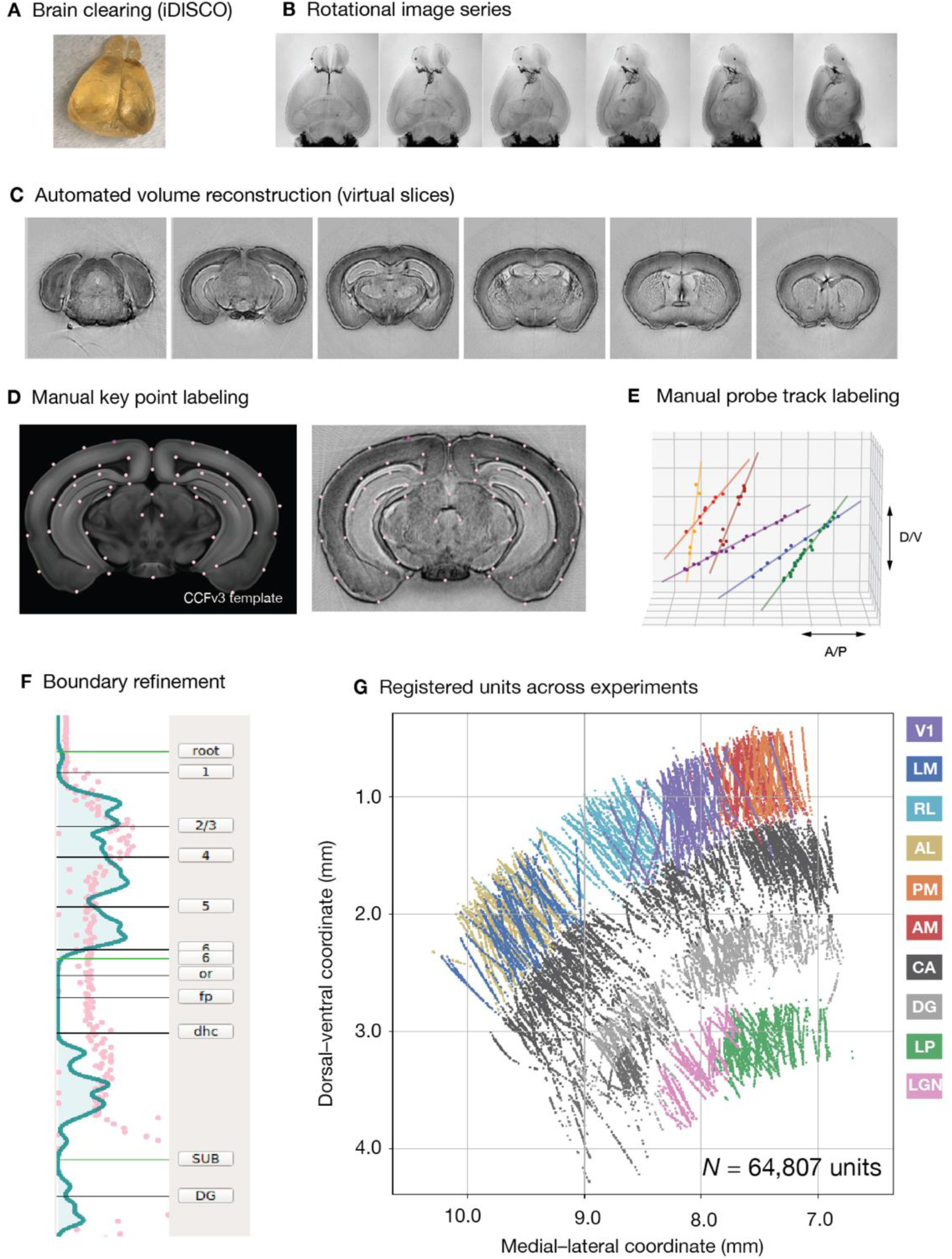
Aligning units with the Common Coordinate Framework (CCFv3). (**A**) Following each experiment, the brain is removed and cleared using a variant of the iDISCO method. (**B**) The cleared brain is imaged at 400 rotational angles using a custom-built optical projection tomography microscope. (**C**) We generated an isotropic 3D volume from rotational images using a computational tomography algorithm. (**D**) Key points from the CCFv3 template brain are manually identified in each individual brain. (**E**) Points along each fluorescently labeled probe track are manually identified in the volume. Using the key points from D, we define a warping function to translate points along the probe axis into the Common Coordinate Framework. (**F**) We then align the regional boundaries to boundaries in the physiological data, primarily the decrease in unit density at the border between cortex and hippocampus, and between hippocampus and thalamus. **(G)** Finally, units in the database are mapped to a 3D location in the CCFv3, and are assigned a structure label.

**Supplementary Figure 6.**
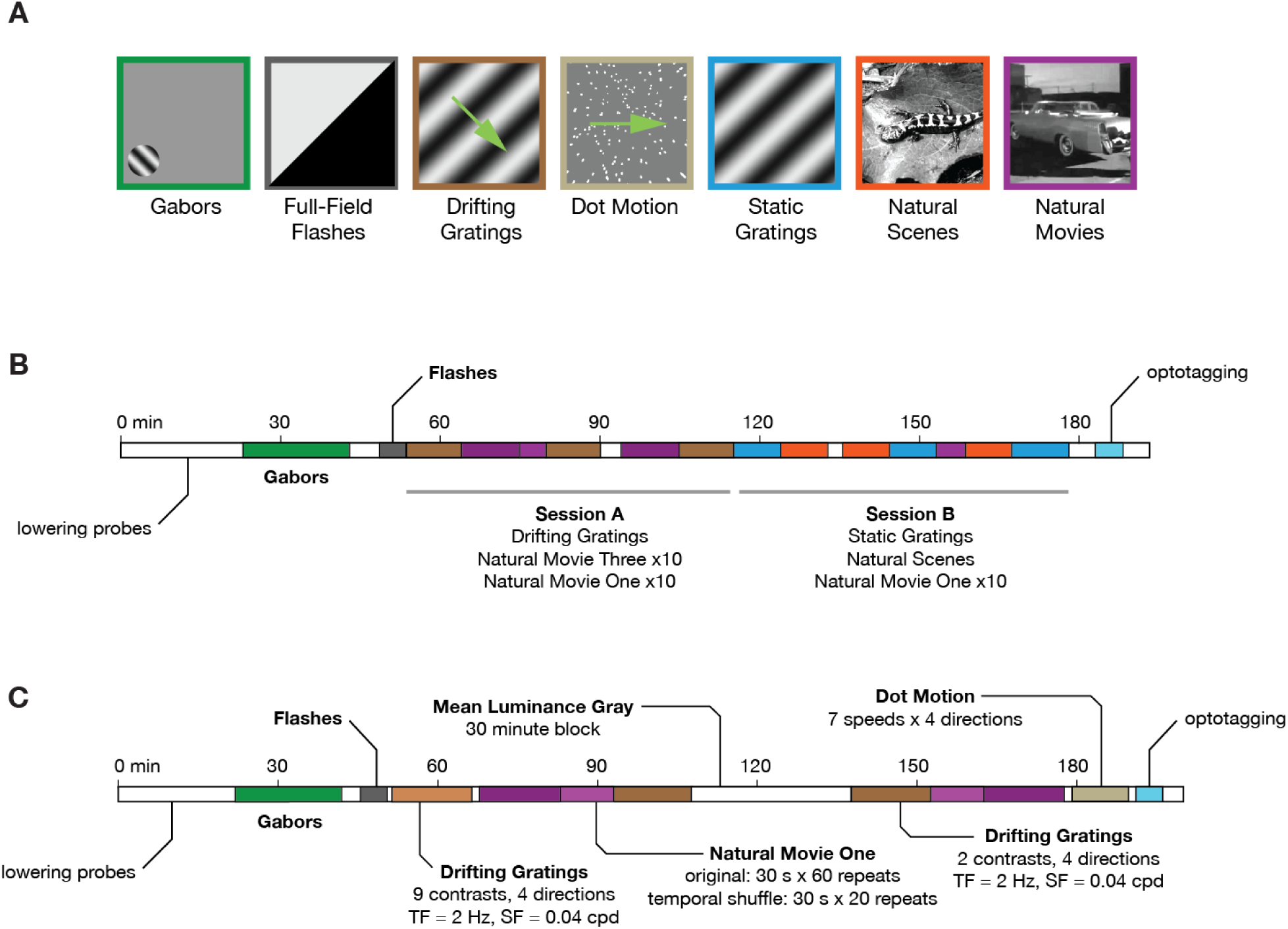
Details of the visual stimulus set. (**A**) Example frames from each type of stimulus. (**B**) Timing diagram for visual stimulus set #1, known as “Brain Observatory 1.1.” **(C)** Timing diagram for visual stimulus set #2, known as “Functional Connectivity.”

**Supplementary Figure 7.**
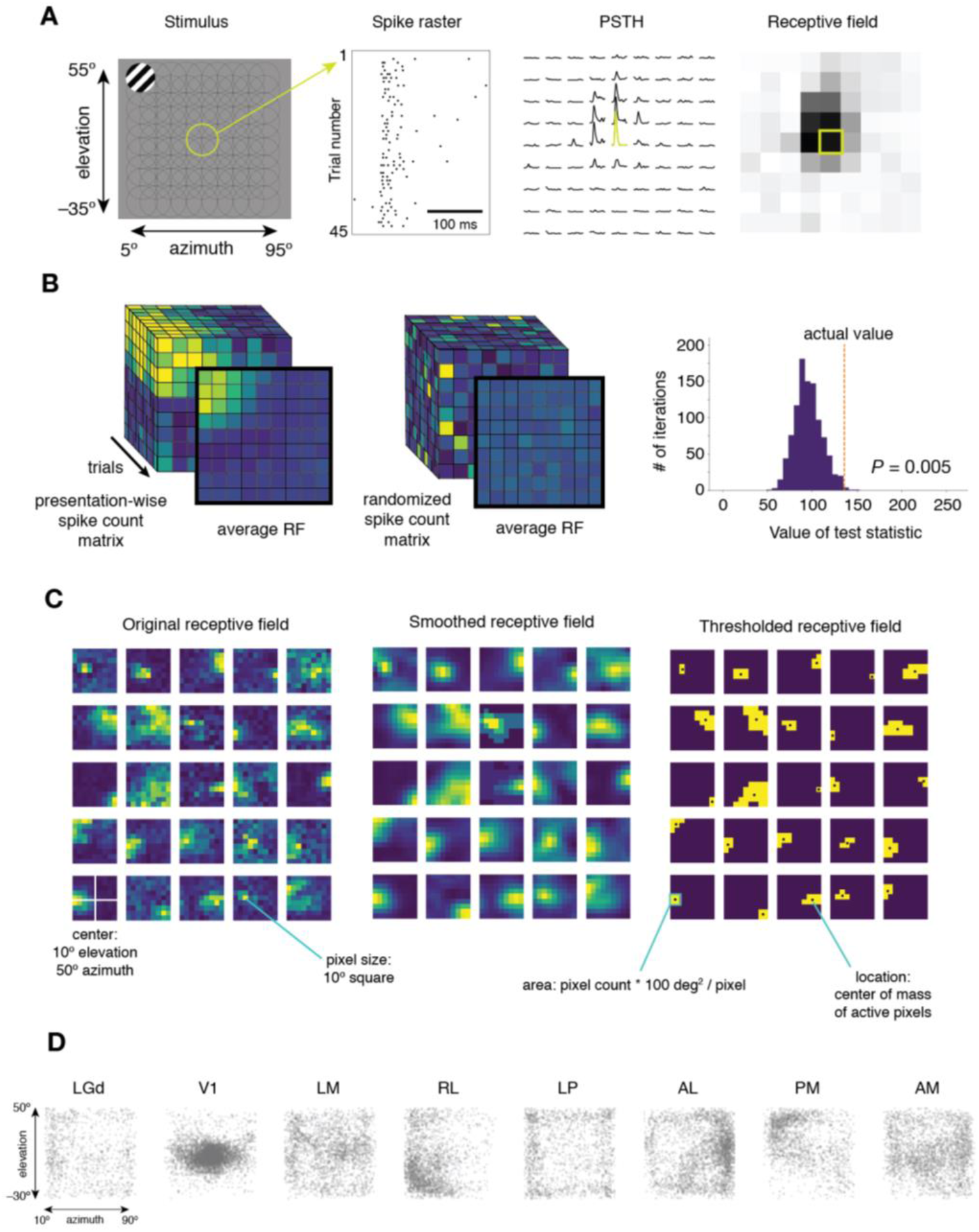
Receptive analysis and unit selection. (**A**) Our receptive field mapping procedure consists of flashing 20° diameter drifting gratings for 250 ms in each of 81 randomized locations on the screen. For each unit, we can construct a spike raster showing the timing of spikes on each of 45 trials with the stimulus at a particular location. Collapsing over trials yields a peri-stimulus time histogram (PSTH) for each location. Collapsing over time yields a spike count for each spatial bin. A matrix of spike counts represents the receptive field for this unit. (**B**) We use a categorical χ² test to determine which units have significant receptive fields. We compare the actual matrix of presentation-wise spike counts (left) to a series of randomized spike count matrices (center) to determine the probability that the receptive field could have occurred by chance (right). (**C**) To calculate receptive field properties, we first smooth the receptive field with a Gaussian filter, then select all pixels above a threshold value. The center of mass of the above-threshold pixels indicates the receptive field location, while the total number of above-threshold pixels indicates the area. These processing steps are shown for 25 receptive fields randomly chosen from one experiment. (**D**) Receptive field locations for all units in our analysis (RF *P*-value < 0.01).

**Supplementary Figure 8.**
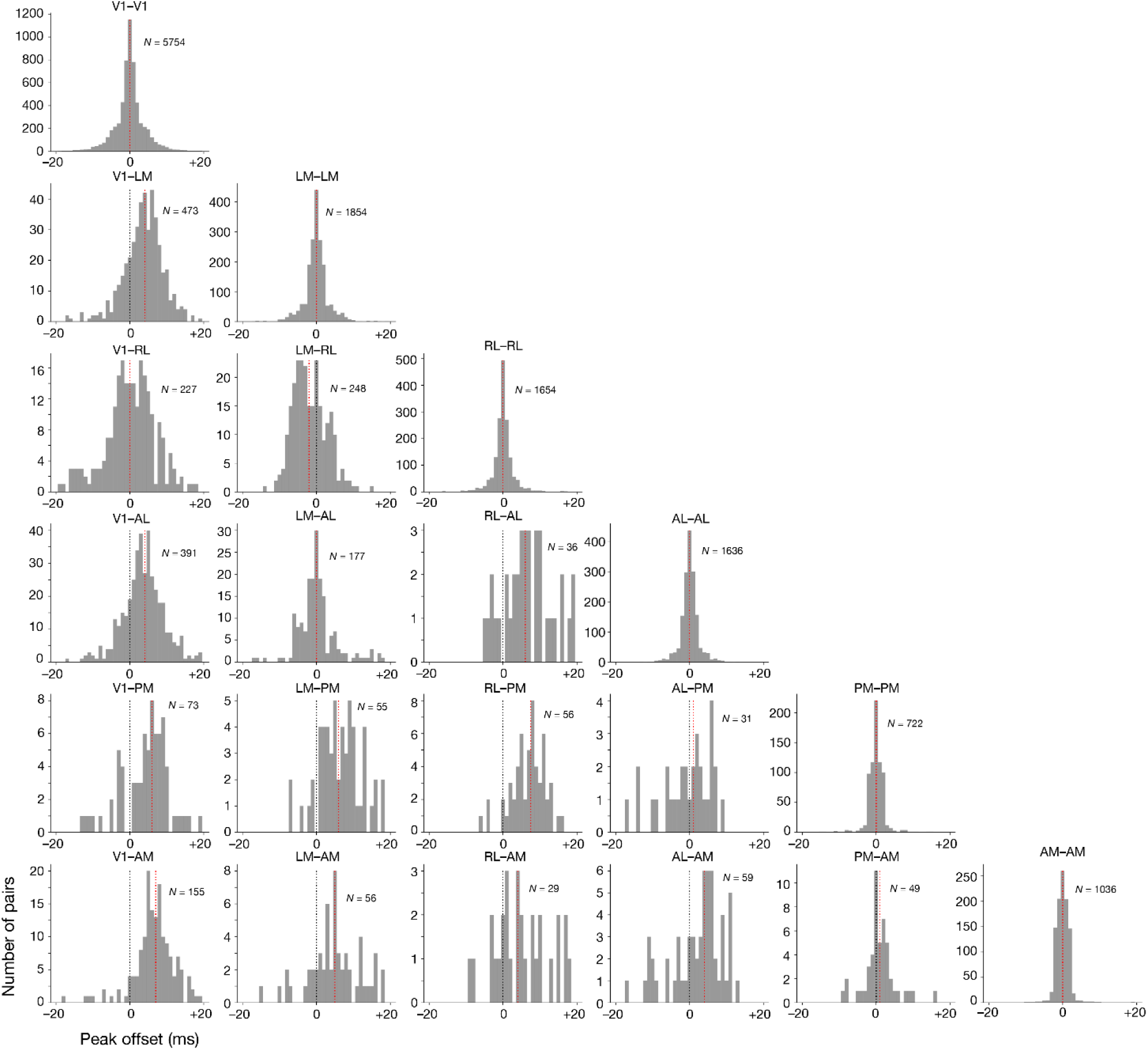
Distributions of pairwise functional delays. Histograms of CCG peak offsets for all pairs of units included in Figure 3.

**Supplementary Figure 9.**
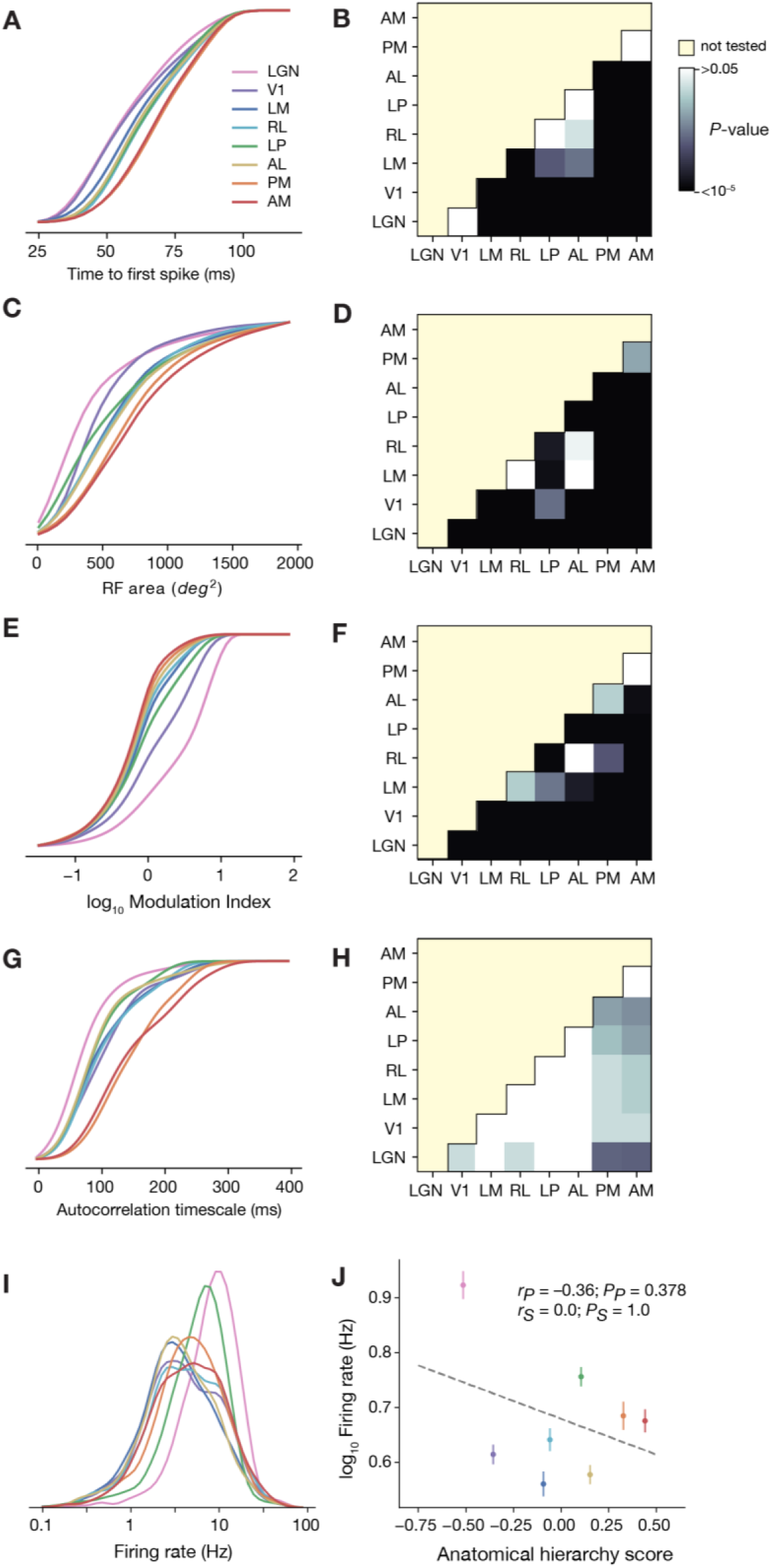
Extended data for Figure 4. **(A)** Cumulative probability function for time to first spike (same data as Figure 4B). **(B)** *P*-values for pairwise comparisons of time to first spike between areas (Wilcoxon rank–sum test with Benjamini-Hochberg False Discovery Rate correction). **(C)** Cumulative probability function for receptive field area (same data as Figure 4E). **(D)** *P*-values for pairwise comparisons of receptive field size between areas. **(E)** Cumulative probability function for modulation index (same data as Figure 4H). **(F)** *P*-values for pairwise comparisons of modulation index between area. **(G)** Cumulative probability function for autocorrelation timescale (same data as Figure 4K). **(H)** *P*-values for pairwise comparisons of autocorrelation timescale between areas. **(I)** Distribution of overall firing rates for all units in each area. (J) Correlation between mean firing rate and anatomical hierarchy score.

**Supplementary Figure 10.**
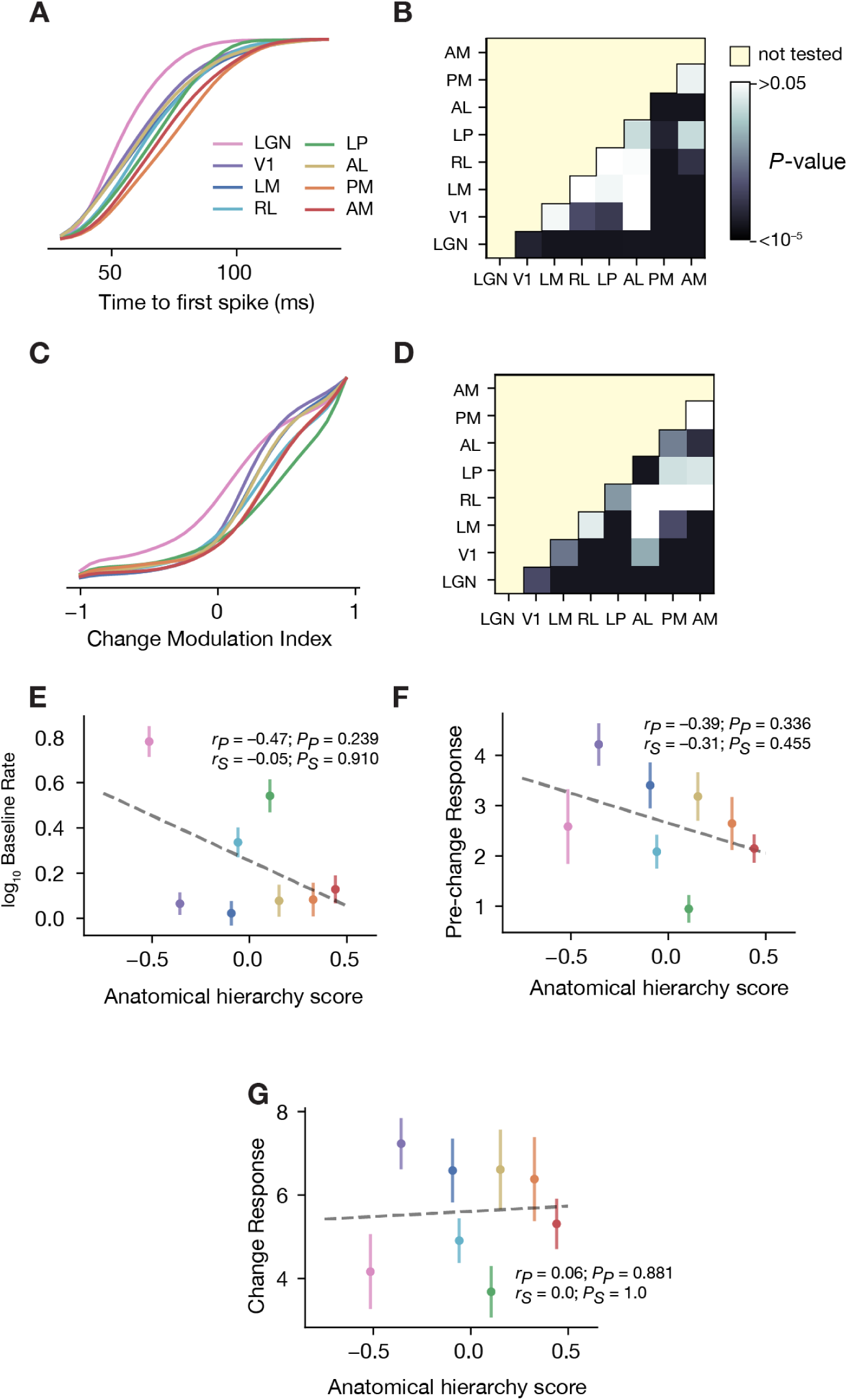
Extended data for Figure 5. **(A)** Cumulative probability function for time to first spike (same data as Figure 5E). **(B)** *P*-values for pairwise comparisons of time to first spike between areas (Wilcoxon rank–sum test with Benjamini-Hochberg False Discovery Rate correction). **(C)** Cumulative probability function for change modulation index (same data as Figure 5F). **(D)** *P*-values for pairwise comparisons of change modulation index between areas **(E)** Correlation between mean baseline firing rate and hierarchy score. **(F)** Correlation between mean pre-change response spike rate and hierarchy score. **(G)** Correlation between mean change response spike rate and hierarchy score.

## Materials and Methods

### 1. Mice

Mice were maintained in the Allen Institute for Brain Science animal facility and used in accordance with protocols approved by the Allen Institute’s Institutional Animal Care and Use Committee.

Wild-type C57BL/6J mice were purchased from Jackson Laboratories at age P25-50. For experiments involving opto-tagging of inhibitory cells, Pvalb-IRES-Cre, Vip-IRES-Cre, and Sst-IRES-Cre mice were bred in-house and crossed with an Ai32 channelrhodopsin reporter line (Madisen et al., 2012). Pvalb-IRES-Cre;Ai32 breeding sets (pairs and trios) consisted of heterozygous Pvalb-IRES-Cre mice crossed with either heterozygous or homozygous Ai32(RCL-ChR2(H134R)_EYFP) mice. Pvalb-IRES-Cre is expressed in the male germline. To avoid germline deletion of the stop codon in the LoxP-STOP-LoxP cassette, Pvalb-IRES-Cre;Ai32 mice were not used as breeders. Sst-IRES-Cre;Ai32 breeding sets (pairs and trios) consisted of heterozygous Sst-IRES-Cre mice crossed with either heterozygous or homozygous Ai32(RCL-ChR2(H134R)_EYFP) mice. Vip-IRES-Cre;Ai32 breeding sets (pairs and trios) consisted of heterozygous Vip-IRES-Cre mice crossed with either heterozygous or homozygous Ai32(RCL-ChR2(H134R)_EYFP) mice. Cre+ cells from Ai32 lines are highly photosensitive, due to expression of Channelrhodopsin-2 (Zhang et al., 2006).

Following surgery, all mice were single-housed and maintained on a reverse 12-hour light cycle. All experiments were performed during the dark cycle. For passive viewing experiments, animals were given ad libitum access to food and water. For behavioral experiments, mice were given an amount of water required to maintain 85% of their initial body weight, with ad libitum access to food.

### 2. Surgery

#### 2.1. Headframe design

To enable co-registration across surgery, intrinsic signal imaging, and electrophysiology rigs, each animal was implanted with a 304 stainless steel headframe that provides access to the brain via a cranial window and permits head fixation in a reproducible configuration (de Vries et al., 2019). The cranial window angle was at 23 degrees of roll and 6 degrees of pitch, referenced to a plane passing through lambda and bregma and the mediolateral axis. Use of this headframe allowed the 5 mm craniotomy to be repeatability centered at *x* = –2.8 mm and *y* = 1.3 mm (origin at lambda).

The headframe was glued to a well made of black acrylic photopolymer that served four functions: (1) shielding the craniotomy and probes during the experiment, (2) providing a surface for precisely aligning the insertion window, (3) routing the animal ground to an exposed gold pin, and (4) holding threads for a plastic cap that protects the craniotomy before and after the experiment.

#### 2.2. Surgical procedures

A pre-operative injection of dexamethasone (3.2 mg/kg, S.C.) was administered 3 h before surgery. Mice were initially anesthetized with 5% isoflurane (1-3 min) and placed in a stereotaxic frame (Model# 1900, Kopf), and isoflurane levels were maintained at 1.5-2.5% for surgery. Body temperature was maintained at 37.5°C. Carprofen was administered for pain management (5-10 mg/kg, S.C.). Atropine was administered to suppress bronchial secretions and regulate hearth rhythm (0.02-0.05 mg/kg, S.C.). An incision was made to remove skin, and the exposed skull was levelled with respect to pitch (bregma-lamda level), roll, and yaw. The headframe was placed on the skull and fixed in place with White C&B Metabond (Parkell). Once the Metabond was dry, the mouse was placed in a custom clamp to position the skull at a rotated angle of 20°, to facilitate creation of the craniotomy over visual cortex. A circular piece of skull 5 mm in diameter was removed, and a durotomy was performed. The brain was covered by a 5 mm diameter circular glass coverslip, with a 1 mm lip extending over the intact skull. The bottom of the coverslip was coated with a layer of silicone to reduce adhesion to the brain surface. The coverslip was secured to the skull with Vetbond (Patterson Veterinary) (Goldey et al., 2014). Kwik-Cast (World Precision Instruments) was added around the coverslip to further seal the implant, and Metabond bridges between the coverslip and the headframe well were created to hold the Kwik-Cast in place. At the end of the procedure, but prior to recovery from anesthesia, the mouse was transferred to a photodocumentation station to capture a spatially registered image of the cranial window (Supplementary Figure 1A).

#### 2.3. Surgery quality control

In cases of excessive bleeding or other complications, the surgical procedure was aborted, and the mouse was euthanized. Mice that completed surgery entered a 7-10 day recovery period that included regular checks for overall health, cranial window clarity, and brain health. If mice failed the first health check, they received another one the following week. Mice that exhibited signs of deteriorating health or damaged brain surface vasculature were not passed on to the next step. Out of 105 mice entering the surgery step, 4 were removed from the pipeline due to QC failures at this stage (Supplementary Figure 2A).

### 3. Intrinsic Signal Imaging

Intrinsic signal imaging (ISI) measures the hemodynamic response of the cortex to visual stimulation across the entire field of view. This technique can be used to obtain retinotopic maps representing the spatial relationship of the visual field (or, in this case, coordinate position on the stimulus monitor) to locations within each cortical area. This mapping procedure was used to delineate functionally defined visual area boundaries to enable targeting of Neuropixels probes to retinotopically defined locations in primary and secondary visual areas (Juavinett et al., 2017).

#### 3.1. Data acquisition

Mice were lightly anesthetized with 1-1.4% isoflurane administered with a SomnoSuite (model #715; Kent Scientific) and vital signs were monitored with a PhysioSuite (model # PS-MSTAT-RT). Eye drops (Lacri99 Lube Lubricant Eye Ointment; Refresh) were applied to maintain hydration and clarity of eyes during anesthesia. Imaging sessions began with a vasculature image acquired under green illumination (527 nm LEDs; Cree Inc., C503B-GCN-CY0C0791). Next the imaging plane was defocused and the hemodynamic response to a visual stimulus was imaged under red light (635 nm LEDs; Avago Technologies, HLMP-EG08-Y2000) with an Andor Zyla 5.5 10 tap sCMOS camera. The stimulus consisted of an alternating checkerboard pattern (20° wide bar, 25° square size) moving across a mean luminance gray background. On each trial, the stimulus bar was swept across the four cardinal axes 10 times in each direction at a rate of 0.1 Hz (Kalatsky and Stryker, 2003). Up to 10 trials were performed on each mouse.

#### 3.2. Data processing

A minimum of three trials were averaged to produce altitude and azimuth phase maps, calculated from the discrete Fourier transform of each pixel. A “sign map” was produced from the phase maps by taking the sine of the angle between the altitude and azimuth map gradients. In the sign maps, each cortical visual area appears as a contiguous red or blue region (Garrett et al., 2014). These maps are used to confirm the cortical area identity of each probe insertion, using the vasculature as fiducial markers (Figure 1C-D, Supplementary Figure 1B).

The altitude and azimuth maps were also used to create a map of eccentricity from the center of visual space (the intersection of 0° altitude and 0° azimuth). Because the actual center of gaze will vary from mouse to mouse, the eccentricity map was shifted to align with the screen coordinates at the center of V1 (which maps to the center of the retina). This V1-aligned eccentricity map was used for probe targeting, to ensure that recorded neurons represent a consistent region on the retina, approximately at the center of the right visual hemifield.

#### 3.3. ISI quality control

The quality control process for the ISI-derived maps included four distinct inspection steps:

3.3.1. The brain surface and vasculature images were inspected post-acquisition for clarity, focus, and position of the cranial window within the field of view.
3.3.2. Individual trials were inspected for visual coverage range and continuity of phase maps, localization of the signal from the amplitude maps and stereotypical organization of sign maps. Only trials respecting these criteria were included in the final average, and a minimum of 3 trials were required.
3.3.3. Visual area boundaries were delineated using automated segmentation, and maps were curated based on stringent criteria to ensure data quality. The automated segmentation and identification of a minimum of six visual areas including V1, LM, RL, AL, AM and PM was required. A maximum of three manual adjustments were permitted to compensate for algorithm inefficiency.
3.3.4. Each processed retinotopic map was inspected for coverage range (35-60° altitude and 60-100° azimuth), bias (absolute value of the difference between max and min of altitude or azimuth range; <10°), alignment of the center of retinotopic eccentricity with the centroid of V1 (<15° apart), and the area size of V1 (>2.8 cm2).

If QC was not passed after the first round of ISI mapping, the procedure was repeated up to two more times to obtain a passing map. In addition to the QC procedures carried out on the ISI-derived maps, the vasculature images were also examined for the presence of white artifacts on the brain surface. White artifacts, an indicator of potential brain damage, were grounds for failing the mouse out of the pipeline. Out of 101 mice entering ISI, 9 did not pass onto habituation due to QC failures during this step (Supplementary Figure 2B).

### 4. Habituation and Behavior Training

#### 4.1. Habituation for passive viewing experiments

Mice underwent two weeks of habituation in sound-attenuated training boxes containing a headframe holder, running wheel, and stimulus monitor (Supplementary Figure 1C). Each mouse was trained by the same operator throughout the 2-week period. During the first week, the operator gently handles the mice, introduces them to the running wheel, and head-fixes them with progressively longer durations each day. During the second week, mice run freely on the wheel and are exposed to visual stimuli for 10 to 50 min per day. The following week, mice undergo habituation sessions of 75 minutes and 100 minutes on the recording rig, in which they view a truncated version of the same stimulus that will be shown during the experiment.

#### 4.2. Behavior training

A subset of mice were trained to perform a change detection task in which one of 8 natural images was continuously flashed (250 ms image presentation followed by 500 ms gray screen) and mice were rewarded for licking when the image identity changed (Figure 5A). The change detection task is described in detail by (Garrett et al., 2019). Briefly, for each trial the time of image change was drawn from an exponential distribution with a minimum of 5 image flashes (3.75 s) and a maximum of 11 flashes (8.25 s). Licking before the image change restarted the trial. Trials in which the mouse licked within 750 ms of image change were “hits,” while licks within 750 ms of non-change catch trials (occurring at the same distribution of times since the last change as change trials) were classified as false alarms (Figure 5B). Mice must perform the task with a *d′* above 1 and have at least 100 contingent (non-aborted) trials for 3 consecutive days prior to moving to the recording rig.

#### 4.3. Habituation quality control

Upon completion of the second week of habituation, mice received an assessment of overall stress levels that reflected observations made by the trainer, including coat appearance, components of the mouse grimace scale, and overall body movements. Out of 92 mice entering habituation for passive viewing experiments, 2 did not pass on to the insertion window implant step (Supplementary Figure 2C)

### 5. Insertion Window Implant

#### 5.1. Window generation

Following the completion of a successful ISI map, a custom insertion window was generated for each mouse. First, six insertion targets were manually drawn on the V1-aligned eccentricity map using a web-based annotation tool. Targets were positioned at the center of retinotopy of V1, LM, AL, AM, and PM; because the retinotopic center of RL often lies on the boundary between RL and S1 barrel cortex, the target location was adjusted to be closer to the geometric center of this area. The coordinates of each target were used to automatically generate the outlines of the insertion window, which was subsequently laser-cut out of 0.5 mm clear PETG plastic (Ponoko). When seated in the headframe well, the window facilitates access to the brain via holes over each of the six visual areas. A solidified agarose/ACSF mixture injected between the brain and the window stabilizes the brain during the recording.

#### 5.2. Surgical procedure

On the day of recording, the cranial coverslip was removed and replaced with an insertion window containing holes aligned to six cortical visual areas. First, the mouse was anesthetized with isoflurane (3%–5% induction and 1.5% maintenance, 100% O_2_) and eyes were protected with ocular lubricant (I Drop, VetPLUS). Body temperature was maintained at 37.5°C (TC-1000 temperature controller, CWE, Incorporated). Metabond bridges were removed from the glass cranial window, followed by the sealing layer of Kwik-Cast. Using a 2 mm silicone suction cup, the cranial window was gently lifted to expose the brain. The insertion window was then placed in the headframe well and sealed with Metabond. An agarose mixture was injected underneath the window and allowed to solidify. The mixture consisted of 0.4 g high EEO Agarose (Sigma-Aldrich), 0.42 g Certified Low-Melt Agarose (Bio Rad), and 20.5 mL ACSF (135.0 mM NaCl, 5.4 mM KCl, 1.0 mM MgCl2, 1.8 mM CaCl2, 5.0 mM HEPES). This mixture was optimized to be firm enough to stabilize the brain with minimal probe drift, but pliable enough to allow the probes to pass through without bending. A layer of silicone oil (30,000 cSt, Aldrich) was added over the holes in the insertion window to prevent the agarose from drying (Supplementary Figure 1D). A 3D-printed plastic cap was screwed into the headframe well to keep out cage debris. At the end of this procedure, mice were returned to their home cages for 1-2 hours.

#### 5.3. Insertion window implant quality control

3 out of 90 mice did not pass through to the recording step due to procedure failures during insertion window implantation. These failures were caused by the headframe coming loose from the skull or excessive bleeding after cranial window removal, after which the mice were euthanized (Supplementary Figure 2D).

### 6. Neuropixels Recordings

#### 6.1. Probes

All neural recordings were carried out with Neuropixels probes (Jun et al., 2017). Each probe contains 960 recording sites, a subset of 374 (“Neuropixels 3a”) or 383 (“Neuropixels 1.0”) of which can be configured for recording at any given time. The electrodes closest to the tip were always used, providing a maximum of 3.84 mm of tissue coverage. The sites are oriented in a checkerboard pattern on a 70 μm wide x 10 mm long shank. Neural signals are routed to an integrated base containing amplification, digitization, and multiplexing circuitry. The signals from each recording site are split in hardware into a spike band (30 kHz sampling rate, 500 Hz highpass filter) and an LFP band (2.5 kHz sampling rate, 1000 Hz lowpass filter). Due to their dense site configuration (20 µm vertical separation along the entire length of the shank), each probe has the capacity to record hundreds of neurons at the same time. Our goal was to insert 6 probes/mouse. Overall, we achieved a penetration success of 5.7 probes/mouse, with failures due to dura regrowth, collisions with the protective cone or opto-tagging fiber optic cable, or probe breakage during manipulation.

The base of each probes contains 32 10-bit analog-to-digital converters (ADCs), each of which are connected to 12 spike-band channels and 12 LFP-band channels via multiplexers. A full cycle of digitization requires 156 samples: 12 samples from each of 12 spike-band channels, and 1 sample from each of 12 LFP-band channels. Each ADC serves a contiguous bank of odd or even channels, so ADC 1 digitizes channels [1,3,5,…,23], ADC 2 digitizes channels [2,4,6,…,24], ADC 3 digitizes channels [25,27,29,…,47], etc. Because of the need for interleaved sampling, common-mode noise will be shared across all channels that are acquired simultaneously, e.g. [1,2,25,26,49,50,…,361,362].

#### 6.2. Experimental rig

The experimental rig (Figure 1B) was designed to allow six Neuropixels probes to penetrate the brain approximately perpendicular to the surface of visual cortex. Each probe is mounted on a 3-axis micromanipulator (New Scale Technologies, Victor, NY), which are in turn mounted on a solid aluminum plate, known as the probe cartridge. The cartridge can be removed from the rig using a pair of pneumatic tool-changers, to facilitate probe replacement and maintenance.

#### 6.3. Workflow Sequencing Engine

The experimental procedure was guided by a Work Sequencing Engine (WSE), a custom GUI written in Python. This software ensured that all experimental steps were carried out in the correct order, reducing trial-to-trial variability and optimizing operator efficiency. The GUI logged the operator ID, mouse ID, and session ID, and ensured that all hardware and software were properly configured. The WSE was also used to start and stop the visual stimulus, the body and eyetracking cameras, and Neuropixels data acquisition.

#### 6.4. Probe alignment

The tip of each probe was aligned to its associated opening in the insertion window using a coordinate transformation obtained via a prior calibration procedure. The XY locations of the six visual area targets were supplied by the Workflow Sequencing Engine (WSE), and these values were translated into XYZ coordinates for each 3-axis manipulator using a custom Python script. The operator then moved each probe into place with a joystick, with the probes fully retracted along the insertion axis.

#### 6.5. Application of CM-DiI

CM-DiI (1 mM in ethanol; ThermoFisher Product #V22888) was used to localize probes during the *ex vivo* imaging step because its fluorescence is maintained after brain clearing, and it has a limited diffusion radius. The probes were coated with CM-DiI before each recording by immersing them one by one into a well filled with dye.

#### 6.6. Head fixation

The mouse was placed on the running wheel and fixed to the headframe clamp with three set screws. Next, the plastic cap was removed from the headframe well and an aluminum cone with 3D-printed wings was lowered to prevent the mouse’s tail from contacting the probes. An IR dichroic mirror was placed in front the right eye to allow the eyetracking camera to operate without interference from the visual stimulus. A black curtain was then lowered over the front of the rig, placing the mice in complete darkness except for the visual stimulus monitor.

#### 6.7. Grounding

A 32 AWG silver wire (A-M Systems) was cemented to the skull during the initial headframe/cranial window surgery. This wire becomes electrically conductive with the brain surface following the application of the ACSF/agarose mixture beneath the insertion window. The wire was pre-soldered to a gold pin embedded in the headframe well, which mates with a second gold pin on the protective cone. The cone pin was soldered to 22 AWG hook-up wire (SparkFun Electronics), which was connected to both the behavior stage and the probe ground. Prior to the experiment, the brain-to-probe ground path was checked using a multimeter.

The reference connection on the Neuropixels probes was permanently soldered to ground using a silver wire, and all recordings were made using an external reference configuration. The headstage grounds (which are contiguous with the Neuropixels probe grounds) were connected together with 36 AWG copper wire (Phoenix Wire). For Neuropixels 3a, two probes had a direct path to animal ground, and the others were wired up serially. All probes were also connected to the main ground via the data cable (a dual coaxial cable). For Neuropixels 1.0, all probes were connected in parallel to animal ground, and were not connected to the main ground through the data cable (a single twisted pair cable).

#### 6.8. Probe insertion

The probe cartridge was initially held approximately 30 cm above the mouse. Once the mouse was secured in the headframe, the cartridge was lowered so the probe tips were approximately 2.5 mm above the brain surface. The probes were then manually lowered one by one to the brain surface until spikes were visible on the electrodes closest to the tip. After the probes penetrated the brain to a depth of ∼100 microns, they were inserted automatically at a rate of 200 µm/min (total of 3.5 mm or less in the brain) to avoid damage caused by rapid insertion (Fiáth et al., 2019). Once the probes reached their targets, they were allowed to settle for 5 to 10 min. Photo-documentation was taken with the probes fully retracted, after the probes reached the brain surface (Supplementary Figure 1E), and again after the probes were fully inserted.

#### 6.9. Data acquisition and synchronization

Neuropixels data was acquired at 30 kHz (spike band) and 2.5 kHz (LFP band) using the Open Ephys GUI (Siegle et al., 2017). Gain settings of 500x and 250x were used for the spike band and LFP band, respectively. Each probe was either connected to a dedicated FPGA streaming data over Ethernet (Neuropixels 3a) or a PXIe card inside a National Instruments chassis (Neuropixels 1.0). Raw neural data was streamed to a compressed format for archiving which was extracted prior to analysis.

Videos of the eye and body were acquired at 30 Hz. The angular velocity of the running wheel was recorded at the time of each stimulus frame, at approximately 60 Hz. Synchronization signals for each frame were acquired by a dedicated computer with a National Instruments card acquiring digital inputs at 100 kHz, which was considered the master clock. A 32-bit digital “barcode” was sent with an Arduino Uno (SparkFun DEV-11021) every 30 s to synchronize all devices with the neural data. Each Neuropixels probe has an independent sample rate between 29,999.90 Hz and 30,000.31 Hz, making it necessary to align the samples offline to achieve precise synchronization. The synchronization procedure used the first matching barcode between each probe and the master clock to determine the clock offset, and the last matching barcode to determine the clock scaling factor. If probe data acquisition was interrupted at any point during the experiment, each contiguous chunk of data was aligned separately. Because one LFP band sample was always acquired after every 12th spike band sample, these data streams could be synchronized automatically once the spike band clock rate has been determined.

To synchronize the visual stimulus to the master clock, a silicon photodiode (PDA36A, Thorlabs) was placed on the stimulus monitor above a “sync square” that flips from black to white every 60 frames. The analog photodiode signal was thresholded and recorded as a digital event by the sync computer. Individual frame times were reconstructed by interpolating between the photodiode on/off events.

#### 6.10. Stimulus monitor

Visual stimuli were generated using custom scripts based on PsychoPy (Peirce, 2007) and were displayed using an ASUS PA248Q LCD monitor, with 1920 x 1200 pixels (21.93 in wide, 60 Hz refresh rate). Stimuli were presented monocularly, and the monitor was positioned 15 cm from the mouse’s right eye and spanned 120° x 95° of visual space prior to stimulus warping. Each monitor was gamma corrected and had a mean luminance of 50 cd/*m*^2^. To account for the close viewing angle of the mouse, a spherical warping was applied to all stimuli to ensure that the apparent size, speed, and spatial frequency were constant across the monitor as seen from the mouse’s perspective.

#### 6.11. Stimuli for passive viewing experiments

All experiments began with a receptive field mapping stimulus consisting of 2 Hz, 0.04 cycles per degree drifting gratings with a 20° circular mask. These Gabor patches randomly appeared at one of 81 locations on the screen (9 x 9 grid) for 250 ms at a time, with no blank interval. The receptive field mapping stimulus was followed by a series of dark or light full-field flashes, lasting 250 ms each and separated by a 2 second inter-trial interval.

Next, mice were shown one of two possible stimulus sets. The first, called “Brain Observatory 1.1” is a concatenation of two sessions from the Two-Photon Imaging Brain Observatory (de Vries et al., 2019) (Supplementary Figure 6B). Drifting gratings were shown with a spatial frequency of 0.04 cycles/deg, 80% contrast, 8 directions (0°, 45°, 90°, 135°, 180°, 225°, 270°, 315°, clockwise from 0° = right-to-left) and 5 temporal frequencies (1, 2, 4, 8, and 15 Hz), with 15 repeats per condition. Static gratings were shown at 6 different orientations (0°, 30°, 60°, 90°, 120°, 150°, clockwise from 0° = vertical), 5 spatial frequencies (0.02, 0.04, 0.08, 0.16, 0.32 cycles/degree), and 4 phases (0, 0.25, 0.5, 0.75); they are presented for 0.25 seconds, with no intervening gray period. The Natural Images stimulus consisted of 118 natural images taken from the Berkeley Segmentation Dataset (Martin et al., 2001), the van Hateren Natural Image Dataset (van Hateren and van der Schaaf, 1998), and the McGill Calibrated Colour Image Database (Olmos and Kingdom, 2004). The images were presented in grayscale and were contrast normalized and resized to 1174 x 918 pixels. The images were presented in a random order for 0.25 seconds each, with no intervening gray period. Two natural movie clips were taken from the opening scene of the movie Touch of Evil (Welles, 1958). Natural Movie One was a 30 second clips repeated 20 times (2 blocks of 10), while Natural Movie Three was a 120 second clip repeated 10 times (2 blocks of 5). All clips were contrast normalized and were presented in grayscale at 30 fps.

The second stimulus set, called “Functional Connectivity,” consisted of a subset of the stimuli from the Brain Observatory 1.1 set shown with a higher number of repeats (Supplementary Figure 6C). Drifting gratings were presented at 4 directions and one temporal frequency (2 Hz) with 75 repeats. A contrast-tuning stimulus consisting of drifting gratings at 4 directions (0°, 45°, 90°, 135°, clockwise from 0° = left-to-right) and 9 contrasts (0.01, 0.02, 0.04, 0.08, 0.13, 0.2, 0.35, 0.6, 1.0) was also shown. The Natural Movie One stimulus was presented a total of 60 times, with an additional 20 repeats of a temporally shuffled version. Last, a dot motion stimulus consisting of approximately 200 1.5° radius white dots on a mean-luminance gray background moving at one of 7 speeds (0°/s, 16°/s, 32°/s, 64°/s, 128°/s, 256°/s, 512°/s) in four different directions (0°, 45°, 90°, 135°, clockwise from 0° = left-to-right) at 90% coherence was shown.

#### 6.12. Stimuli for behavioral experiments

Mice carried out one hour of a change detection task as described in (Garrett et al., 2019). Following the behavior session, the lick spout was retracted and receptive field mapping stimuli and full-field flashes were presented for 25 minutes, with the same parameters as those used in the passive viewing experiments. All other aspects of the rig, including the running wheel, stimulus monitor, and electrophysiological recordings were the same as for the passive viewing experiments.

#### 6.13. Probe removal and cleaning

Once the stimulus set was over, probes were retracted from the brain at a rate of 1 mm/s, after which the probe cartridge was raised to its full height. The protective cap was screwed into the headframe well, then mice were removed from head fixation and returned to their home cages overnight. Probes were immersed in a well of 1% Tergazyme for ∼24 hours, which was sufficient to remove tissue and silicone oil prior to the next recording session.

#### 6.14. Quality control for the Neuropixels recording session

Neuropixels recording sessions were subjected to the following QC criteria (Supplementary Figure 2E):

##### 6.14.1. Eye foam

If white buildup around the eye obscures the pupil, the experiment is cancelled and the session is failed (8 mice).

##### 6.14.2. Bleeding

If bleeding resulting from the window implant or the probe insertion obscures the vasculature, the session is failed (4 mice).

##### 6.14.3 Probe insertion

If fewer than four probes successfully enter the brain, the session is failed (1 mouse).

##### 6.14.4. Dropped frames

If the stimulus monitor photodiode measures >60 delayed frames, the session is failed (1 mouse).

##### 6.14.5. Missing files

If any critical files are overwritten, the session is failed (2 mice).

##### 6.14.6. Noise levels

If high RMS noise levels in the spike band persist after median subtraction, the session is failed (4 mice).

##### 6.14.7. Probe drift

If one or more probes exhibit >80 microns of drift over the course of the experiment, the session is failed (6 mice). Typical drift levels are around 40 microns, and drift levels are highly correlated across probes.

In total, out of 87 mice entering the recording step, 61 passed session-level QC.

### 7. *Ex Vivo* Imaging

#### 7.1. Tissue clearing

Mice were perfused with 4% paraformaldehyde (PFA) (after induction with 5% isoflurane and 1 L/min of O2). The brains were preserved in 4% PFA, rinsed with 1x phosphate buffered saline (PBS) the next morning, and stored at 4°C in PBS. Next, brains were run through a tissue clearing process based on the iDISCO method (Renier et al., 2014). This procedure uses different solvents which dehydrate and delipidate the tissue. The first day, the brains were immersed in different concentrations of methanol (20, 40, 60) for an hour each, then overnight in 80% methanol. On the second day, they were dipped into 100% methanol (twice for one hour) and then into a mixture of 1/3 methanol and 2/3 dichloromethane overnight. On the third day, the brains were moved from pure dichloromethane (2 x 20 min) to pure dibenzyl ether, where they remained for 2 days until clearing was complete (Supplementary Figure 5A).

#### 7.2. Optical projection tomography (OPT)

Whole-brain 3D imaging was accomplished with optical projection tomography (OPT) (Nguyen et al., 2017; Sharpe, 2002; Wong et al., 2013). The OPT instrument consisted of collimated light sources for transmitted illumination (on-axis white LED, Thorlabs MNWHL4 with Thorlabs SM2F32-A lens and Thorlabs DG20-600 diffuser) or fluorescence excitation (off-axis Thorlabs M530L3, with Thorlabs ACL2520U-DG6-A lens and Chroma ET535/70m-2P diffuser), a 0.5x telecentric lens (Edmund Optics 62-932) with emission filter (575 nm LP, Edmund Optics 64-635), and a camera (IDS UI-3280CP). The specimen was mounted on a rotating magnetic chuck attached to a stepper motor, which positioned the specimen on the optical axis and within a glass cuvette filled with dibenzyl ether. The stepper motor and illumination triggering were controlled with an Arduino Uno (SparkFun DEV-11021) and custom shield including a Big Easy Driver (SparkFun ROB-12859). Instrument communication and image capture was accomplished with MicroManager (Edelstein et al., 2014).

A series of 400 images were captured with transmitted LED illumination with each image captured with the specimen rotated 0.9 degrees relative to the previous position. This series of 400 images was repeated with the fluorescence excitation LED. Each channel was stored as a separate OME-TIFF dataset before extracting individual planes and metadata required for reconstruction using a custom Python script (Supplementary Figure 5B).

Isotropic 3D volumes were reconstructed from these projection images using NRecon (Bruker). The rotation axis offset and region-of-interest bounds were set for each image series pair using the transmitted channel dataset, then the same values applied to the fluorescence channel dataset. A smoothing level of 3 using a Gaussian kernel was applied to all images. Reconstructions were exported as single-plane 16-bit TIFF images taken along the rotation axis with final voxel size of 7.9 µm per side (Supplementary Figure 5C).

#### 7.3. Registering Probes to the Common Coordinate Framework

Reconstructed brains were downsampled to 10 µm per voxel and roughly aligned to the Allen Institute Common Coordinate Framework (CCFv3) template brain using an affine transform. The volume was then cropped to a size of 1023 x 1024 x 1024 and converted to Drishti format (http://sf.anu.edu.au/Vizlab/drishti). Next, 6-54 registration points were marked in up to 14 coronal slices of the individual brain by comparing to the CCFv3 template brain, obtained from (Shamash et al., 2018) (Supplementary Figure 5D). Fluorescent probe tracks were manually labeled in coronal slices of the individual brain, and the best-fit line was found using singular value decomposition (Supplementary Figure 5E). The registration points were used to define a 3D nonlinear transform (VTK thinPlateSplineTransform), which was used translate each point along the probe track into the CCFv3 coordinate space. Each CCFv3 coordinate corresponds to a unique brain region, identified by its structure acronym (e.g., CA3, LP, VISp, etc.). A list of CCFv3 structure acronyms along each track was compared to the physiological features measured by each probe (e.g., unit density, LFP theta power; Supplementary Figure 5F). The location of major structural boundaries were manually adjusted to align the CCFv3 labels with the physiology data, and each recording channel (and its associated units) was assigned to a unique CCFv3 structure (Supplementary Figure 5G). White matter structures were not included; any units mapped to a white matter structure inherited the gray matter structure label that was immediately ventral along the probe axis.

#### 7.4. Identification of cortical visual area targets

To confirm the identity of the cortical visual areas, images of the probes taken during the experiment were compared to images of the brain surface vasculature taken during the ISI session. Vasculature patterns were used to overlay the visual area map on an image of the brain surface with the probes inserted. When done in custom software, key points were selected along the vasculature on both images and a perspective transform (OpenCV) was performed to warp the insertion image to the retinotopic map. When done manually, the overlap of both images was done in Photoshop or Illustrator (Adobe Suite). In both cases, the probe entry points were manually annotated. Finally, an area was assigned to each probe. Overall, successful targeting of the 6 target visual areas occurred at the following rates: 89% for AM, 72% for PM, 98% for V1, 85% for LM, 79% for AL, and 90% for RL. A small subset of penetrations were mapped to LI, MMA, or MMP (Zhuang et al., 2017). Penetration points that could not be unambiguously associated with a particular visual area were classified as “VIS.” If the cortical area label obtained via CCFv3 registration did not match the area identified in the insertion image overlay, the insertion image overlay took precedence.

#### 7.5. *Ex vivo* imaging quality control

**7.6.**

QC was performed on a probe-by-probe, rather than a mouse-by-mouse basis. Some probes were not visible in the OPT images due to faint CM-DiI signal or reconstruction artifacts caused by air bubbles in the tissue (Supplementary Figure 2F). In total, 284 out of 332 probes were mapped to the CCFv3. Probes that failed the *ex vivo* imaging step were not excluded from further analysis, but only included structure labels for channels in cortex (with the bottom of cortex identified based on the drop in unit density between cortex and hippocampus).

### 8. Spike Sorting

#### 8.1. Data pre-processing

Data was written to disk in a format containing the original 10-bit samples from each ADC. These files were backed up to a tape drive, then extracted to a new set of files that represent each sample as a 16-bit integer, scaled to account for the gain settings on each channel. Separate data files were generated for the LFP band and spike band, along with additional files containing the times of synchronization events. The extracted files consume approximately 36% more disk space than the originals.

Prior to spike sorting, the spike-band data passed through 4 steps: offset removal, median subtraction, filtering, and whitening. First, the median value of each channel was subtracted to center the signals around zero. Next, the median across channels was subtracted to remove common-mode noise. While Neuropixels have been measured to have a spike-band RMS noise levels of 5.1 µV in saline (Jun et al., 2017), this cannot be achieved in practice when recording in vivo. The signals become contaminated by background noise in neural tissue; movement artifacts associated with animal locomotion, whisking, and grooming; and electrical noise introduced by the additional wiring required to support multiple probes on one rig. To remove noise sources that are shared across channels, the median was calculated across channels that are sampled simultaneously, leaving out adjacent (even/odd) channels that are likely measuring the same spike waveforms, as well as reference channels that contain no signal. For each sample, the median value of channels N:24:384, where N = [1,2,3,…,24], was calculated, and this value was subtracted from the same set of channels. This method rejects high-frequency noise more effectively than subtracting the median of all channels, at the cost of leaving a residual of ∼2 µV for large spikes, visible in the mean waveforms. Given that this value is well below the RMS noise level of the Neuropixels probes under ideal conditions, it should not affect spike sorting. The original data is over-written with the median-subtracted version, with the median value of each block of 16 channels saved separately, to allow reconstruction of the original signal if necessary. The median-subtracted data file is sent to the Kilosort2 Matlab package (https://github.com/mouseland/kilosort2, commit 2fba667359dbddbb0e52e67fa848f197e44cf5ef, April 8, 2019), which applies a 150 Hz high-pass filter, followed by whitening in blocks of 32 channels. The filtered, whitened data is saved to a separate file for the spike sorting step.

#### 8.2. Kilosort2

Kilosort2 was used to identify spike times and assign spikes to individual units (Stringer et al., 2019). Traditional spike sorting techniques extract snippets of the original signal and perform a clustering operation after projecting these snippets into a lower-dimensional feature space. In contrast, Kilosort2 attempts to model the complete dataset as a sum of spike “templates”. The shape and locations of each template is iteratively refined until the data can be accurately reconstructed from a set of *N* templates at *M* spike times, with each individual template scaled by an amplitude, *a*. A critical feature of Kilosort2 is that it allows templates to change their shape over time, to account for the motion of neurons relative to the probe over the course of the experiment. Stabilizing the brain using an agarose-filled plastic window has virtually eliminated probe motion associated with animal running, but slow drift of the probe over ∼3-hour experiments is still observed. Kilosort2 is able to accurately track units as they move along the probe axis, eliminating the need for the manual merging step that was required with the original version of Kilosort (Pachitariu et al., 2016). The spike-sorting step runs in approximately real time (∼3 hours per session) using a dual-processor Intel 4-core, 2.6 GHz workstation with an NVIDIA GTX 1070 GPU.

#### 8.3. Removing putative double-counted spikes

The Kilosort2 algorithm will occasionally fit a template to the residual left behind after another template has been subtracted from the original data, resulting in double-counted spikes. This can create the appearance of an artificially high number of ISI violations for one unit or artificially high zero-time-lag synchrony between nearby units. To eliminate the possibility that this artificial synchrony will contaminate data analysis, the outputs of Kilosort2 are post-processed to remove spikes with peak times within 5 samples (0.16 ms) and peak waveforms within 5 channels (∼50 microns). This process removes >10 within-unit overlapping spikes from 2.5 ± 1.8% of units per session. It removes 2.05 ± 0.65% of spikes in total, after accounting for between-unit overlapping spikes.

#### 8.4. Removing units with artifactual waveforms

Kilosort2 generates templates of a fixed length (2 ms) that matches the time course of an extracellularly detected spike waveform. However, there are no constraints on template shape, which means that the algorithm often fits templates to voltage fluctuations with characteristics that could not physically result from the current flow associated with an action potential. The units associated with these templates are considered “noise,” and are automatically filtered out based on 3 criteria: spread (single channel, or >25 channels), shape (no peak and trough, based on wavelet decomposition), or multiple spatial peaks (waveforms are non-localized along the probe axis). The automated algorithm removed 94% of noise units, or 26% of total units. A final manual inspection step was used to remove an additional 2140 noise units across all experiments (Supplementary Figure 3).

#### 8.5. Spike sorting quality control

All units not classified as noise are packaged into Neurodata Without Borders (NWB) files for potential further analysis. Because different analyses may require different quality thresholds for defining inclusion criteria, we calculate a variety of metrics that can be used to filter units. These metrics are based on both the physical characteristics of the units’ waveforms, or their isolation with respect to other units from the same recording (Supplementary Figure 4A).

##### 8.5.1. Firing rate

*N*/*T*, where *N* = number of spikes in the complete session and *T* = total time of the recording session in seconds.

##### 8.5.2. Presence ratio

The session was divided into 100 equal-sized blocks; the presence ratio is defined as the fraction of blocks that include 1 or more spikes from a particular unit. Units with a low presence ratio are likely to have drifted out of the recording, or could not be tracked by Kilosort2 for the duration of the experiment.

##### 8.5.3. Maximum drift

To compute the maximum drift for one unit, the peak channel was calculated from the top principal components of every spike. Next, the peak channel values are binned in 51 s intervals, and the median value is calculated across all spikes in each bin (assuming at least 10 spikes per bin). The maximum drift is defined as the difference between the maximum peak channel and the minimum peak channel across all bins. The average maximum drift across all units is used to identify sessions with a high amount of probe motion relative to the brain.

##### 8.5.4. Waveform amplitude

The difference (in microvolts) between the peak and trough of the waveform on a single channel.

##### 8.5.5. Waveform spread

Spatial extent (in microns) of channels where the waveform amplitude exceeds 12% of the peak amplitude.

##### 8.5.6. Waveform duration

Difference (in ms) of the time of the waveform peak and trough on the channel with maximum amplitude.

##### 8.5.7. Inter-spike-interval (ISI) violations

This metric searches for refractory period violations that indicate a unit contains spikes from multiple neurons. The ISI violations metric represents the relative firing rate of contaminating spikes. It is calculated by counting the number of violations <1.5 ms, dividing by the amount of time for potential violations surrounding each spike, and normalizing by the overall spike rate. It is always positive (or 0), but has no upper bound. See (Hill et al., 2011) for more details.

##### 8.5.8. Signal-to-noise ratio (SNR)

After selecting 1000 individual spike waveforms on the channel with maximum amplitude, the mean waveform on that channel was subtracted. SNR is defined the ratio between the waveform amplitude and 2x the standard deviation of the residual waveforms (Suner et al., 2005). Because this definition of SNR assumes that waveforms remain stable over time, changes in a unit’s waveform as a result of probe motion will cause this metric to be inaccurate. In addition, because it is only calculated for the peak channel, this metric does not necessarily reflect the overall isolation quality of a unit when taking into account all available information.

##### 8.5.9. Isolation distance

The square of the Mahalanobis distance required to find the same number of “other” spikes as the total number of spikes for the unit in principal component space (Schmitzer-Torbert et al., 2005). Similarly to SNR, isolation distance is not tolerant to electrode drift, and changes in waveform shape over time can reduce the isolation distance calculated over the entire session.

##### 8.5.10. d′

Linear discriminant analysis is used to find the line of maximum separation in PC space. *d′* indicates the separability of the unit of interest from all other units. See (Hill et al., 2011) for more information. This metric is not tolerant to electrode drift, and changes in waveform shape over time can reduce the value of *d′* calculated over the entire session.

##### 8.5.11. Amplitude cutoff

This metric provides an approximation of a unit’s false negative rate. First, a histogram of spike amplitudes is created, and the height of the histogram at the minimum amplitude is extracted. The percentage of spikes above the equivalent amplitude on the opposite side of the histogram peak is then calculated. If the minimum amplitude is equivalent to the histogram peak, the amplitude cutoff is set to 0.5 (indicating a high likelihood that >50% of spikes are missing). This metric assumes a symmetrical distribution of amplitudes and no drift, so it will not necessarily reflect the true false negative rate.

##### 8.5.12. Nearest neighbors hit rate

For each spike belonging to the unit of interest, the four nearest spikes in principal-component space are identified. The “hit rate” is defined as the fraction of these spikes that belong to the unit of interest. This metric is based on the “isolation” metric from (Chung et al., 2017). Again, electrode drift that alters waveform shape can negatively impact this metric without necessarily changing the isolation quality of a unit at any given timepoint.

Filtering of units based on quality metrics and other criteria is illustrated in Supplementary Figure 4B.

### 9. Data analysis

#### 9.1. Receptive field analysis

The receptive field for one unit is defined as the 2D histogram of spike counts at each of 81 locations of the Gabor stimulus (9 x 9 pixels, 10° separation between pixel centers, Supplementary Figure 7A).

A chi-square test for independence was used to assess the presence of a significant receptive field. A chi-square test statistic was computed 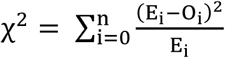, where 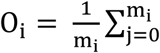 R_i,j_ is the observed average response (R) of the unit over m presentations of the Gabor stimulus at location i, and 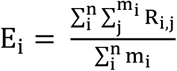 is the expected (grand average) response per stimulus presentation. A *P*-value was then calculated for each unit by comparing the test statistic against a null distribution of 1,000 test statistics, each computed from the unit’s responses after shuffling the locations across all presentations (Supplementary Figure 7B).

To compute the receptive field area and center location, each receptive field was first smoothed using a Gaussian filter (sigma = 1.0). The smoothed receptive field was thresholded at max(RF) – std(RF), a value that provided good agreement with the qualitative receptive field boundaries. The receptive field center location was calculated based on the center of mass of the largest contiguous area above threshold, and its area was equivalent to its pixel-wise area multiplied by 100 degrees^2^ (Supplementary Figure 7C).

#### 9.2. Cross-correlation analysis

To measure the functional interactions between pairs of units, cross-correlograms (CCGs) were used (Gerstein and Perkel, 1972; Jia et al., 2013; Smith and Kohn, 2008). The CCG is defined as:

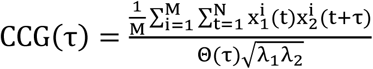

where *M* is the number of trials, *N* is the number of bins in the trial, 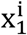 and 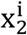 are the spike trains of the two units on trial i, τ is the time lag relative to reference spikes, and λ_1_and λ_2_ are the mean firing rates of the two units. The CCG is essentially a sliding dot product between two spike trains. θ(τ) is the triangular function which corrects for the overlap time bins caused by the sliding window. To correct for firing rate dependency, we normalized the CCG by the geometric mean spike rate. An individually normalized CCG is computed separately for each drifting grating orientation and averaged across orientation to obtain the CCG for each pair of units.

A jitter correction method (Harrison and Geman, 2009; Smith and Kohn, 2008) was used to remove stimulus-locked correlations and slow temporal correlations from the original CCG.

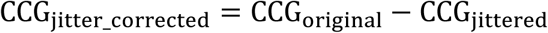

The jitter corrected CCG is created by subtracting the CCG calculated from a jittered spike train where spike times within a small time window are randomly shuffled across trials within that window. For our measurement, a 25 ms jitter window was chosen based on previous studies (Jia et al., 2013; Zandvakili and Kohn, 2015).

The jitter-corrected CCG was deemed to be significant sharp peak if the CCG peak occured within a 10 ms time lag and the magnitude of CCG peak was at least 7-fold larger than the standard deviation of the CCG flanks (±50 to 100 ms).

A Wilcoxon rank sum test was used to compare the distribution of CCG peak offsets between neighboring areas (defined by the anatomical hierarchical score) and the distribution of CCG peak offset within an area. The significance test was performed within each mouse, and the *P*-values were combined across 25 mice using Fisher’s method. V1–LM vs. V1–V1, *P* = 0; LM– RL vs. LM–RL, *P* = 1.9e-5; RL–AL vs. RL–RL, *P* = 2.4e-5; AL–PM vs. AL–AL, *P* = 0.081; PM–AM vs. PM–PM, *P* = 3.2e-4. All between-area distributions are significantly different from the within-area distributions at the 5% confidence level, except for AL–PM.

#### 9.3. Response latency

Response latency is calculated as the time to first spike (TTF). TTF is estimated in each trial by looking for the time of first spike 30 ms after stimulus onset. If no spike is detected within 250 ms after stimulus onset, that trial is not included. The overall latency for each unit is defined as the median TTF across trials.

#### 9.4. Modulation index

The stimulus modulation index reflects how spiking activity of each unit is modulated by the temporal frequency of the drifting grating stimulus (Matteucci et al., 2019; Wypych et al., 2012). It is defined as:

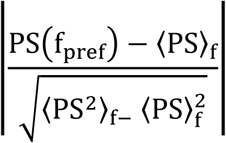

where PS indicates the power spectral density of the peristimulus time histogram (PSTH), and denotes the averaged power over all frequencies; f_pref_ is the preferred temporal frequency of the unit. This metric quantifies the difference between spiking response power at each unit’s preferred frequency and the total power. The power spectrum was computed using Welch’s method on the 10 ms-binned PSTH for each unit’s preferred condition.

#### 9.5. Autocorrelation timescale

We calculated the autocorrelation for each unit during the 250 ms presentation period of the full-field flash stimulus. We estimated autocorrelation timescale in each mouse by calculating a mean of autocorrelation across units within each area, and then fitting an exponential decay to estimate the timescale.

#### 9.6. Analysis of neural responses during the change detection task

For each unit, spike density functions (SDFs) were calculated by convolving spike times relative to each image change or the image flash preceding image change (“pre-change”) with a causal exponential filter (decay time constant = 5 ms). The firing rate during a baseline window 250 ms immediately preceding each image change or pre-change flash was subtracted from each SDF. Mean SDFs were then calculated by averaging across all image change or pre-change flashes. Units were included in further analysis if their mean firing rate was greater than 0.1 spikes/s and the peak of the mean SDF (during a response window from 30 to 280 ms following image change) was greater than 5 times the standard deviation of the mean SDF during the baseline window.

Responses to image change and pre-change were calculated as the mean baseline-subtracted firing rate during the response window. We defined the change modulation index for each unit as the difference between the mean response to image change and pre-change divided by their sum (Figure 5D).

#### 9.7. Eye and pupil tracking

A single, universal eye tracking model was trained in DeepLabCut (Mathis et al., 2018), a ResNET-50 based network, to recognize up to twelve tracking points each around the perimeter of the eye, the pupil, and the corneal reflection. A published numerical routine (Halir and Flusser, 1998) was used to fit ellipses to each set of tracking points. For each ellipse, the following parameters were calculated: center coordinates, half-axes, and rotation angle. Fits were performed on each frame if there at least six tracked points and a confidence of l > 0.8 as reported by the output of DeepLabCut. For frame where there were less than 6 tracked points above the confidence threshold, the ellipse parameters were set to not-a-number (NaN).

The training data set contained two sources of hand-annotated data: (1) 3 frames from each of 40 randomly selected movies. On each frame, 8 points from were annotated around the eye and pupil. The center of the corneal reflection was annotated with a single point. (2) 4150 frames with the pupil and corneal reflections annotated with ellipses.

#### 9.8. Anatomical hierarchy analysis

A detailed description of the unsupervised construction of a data-driven anatomical hierarchy is available in (Harris et al., 2019). Here we provide a summary of how the anatomical hierarchy of the six visual cortical areas (V1, LM, AL, RL, PM, AM) and two thalamic nuclei (LGN, LP) was constructed based on the anatomical connectivity. Specifically, the anatomical hierarchy was uncovered based on cortical lamination patterns of the structural connections among the cortical and thalamic regions of interest, obtained from Cre-dependent viral tracing experiments.

To classify laminar patterns of cortico-cortical (CC) and thalamo-cortical (TC) connections and to assign a direction to each cluster of laminar patterns, we used a large-scale dataset on cell class-specific connectivity among all 37 cortical areas and 24 thalamic nuclei defined using 15 Cre driver transgenic lines (849 cortical and 81 thalamic experiments; 7063 unique source-target-Cre line combinations), available in Harris et al (2019). For each transgenic line, the strength and layer termination pattern of the connections were quantified based on *relative layer density*, the fraction of the total projection signal in each layer scaled by the relative layer volumes in that target. For the connections above a threshold (10^−1.5^), unsupervised clustering of the layer termination patterns was performed, yielding nine clusters of distinct cortical layer termination patterns of CC and TC connections. See Figure 5A,B of Harris et al (2019) for a schematic of the nine types of cortical target lamination patterns.

Following the classification of the nine clusters of the laminar patterns, an unsupervised method was employed to simultaneously assign a direction to a cluster type and to construct a hierarchy by maximizing the self-consistency of the obtained hierarchy. The mapping function *M*_*CC*_ maps a type of CC connection cluster (*C*_*Ti*,*j*_ ∈ {1, … 9}, where *C*_*Ti*,*j*_ denotes the layer termination pattern of the connection from area *j* to area *i* for Cre-line *T*) to either feedforward (*M*_*CC*_= 1) or feedback (*M*_*CC*_ = −1) type, i.e., *M*_*CC*_: {1, …, 9} → {−1,1}. Similarly, the mapping function *M*_*TC*_ of the thalamocortical layer termination types to either direction is defined as *M*_*TC*_: {1, …, 9} → {−1,1}. By constructing the hierarchy of all 37 cortical areas and 24 thalamic nuclei, Harris et al (2019) found the optimal mapping function that maximizes the self-consistency measured by the *global hierarchy score* (Refer to Eq 5 and Eq 10 of Harris et al (2019) to see how the global hierarchy score was defined for CC and TC connections, respectively.). Specifically, the optimal mapping for CC connections assigns connections of cluster 2,6, and 9 to one direction (feedback) and 1,3,4,5,7, and 8 to the opposite direction (feedforward). For TC connections, the most self-consistent hierarchy that maximizes the global hierarchy score is obtained when connections of cluster 2 and 6 correspond to feedback and the rest to feedforward patterns (Figure 6A of Harris et al (2019)).

With these mapping functions *M*_*CC*_ and *M*_*TC*_ obtained from the construction of the all-area hierarchy (Figure 6A of Harris et al (2019)), the hierarchical organization of the six visual cortical areas (V1, LM, AL, RL, PM, AM) and the two thalamic nuclei (LGN, LP) was constructed using only the connections among these 8 regions. We first uncovered the cortical hierarchy using the intra-cortical connections among the six cortical areas: V1, LM, AL, RL, PM, and AM (240 unique “source-target-Cre line” combinations). The initial hierarchical position of a cortical area is defined as:

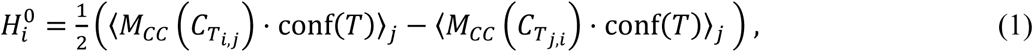

where the first term describes the average direction of connections to area *i*, and thus represents the hierarchical position of the area as a target. The second term on the other hand, represents the average direction of connections from area *i*, depicting the hierarchical position of the area as a source. To account for the Cre-line-specific bias, the Cre-dependent confidence measure, 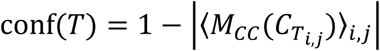 is included. The initial hierarchy score 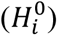 of each area *i* then is iterated using a two-step iterative scheme until the fixed point is reached:

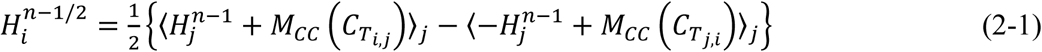

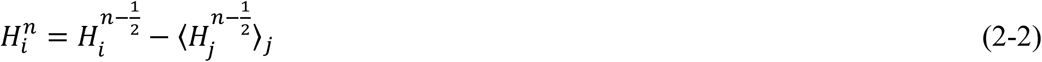

where *n* refers to iterative steps.

After hierarchical positions of cortical areas are found based on CC connections, the hierarchical positions of LGN and LP relative to the cortical areas were computed by including TC connections from LGN and LP to the six visual cortical areas (25 unique “source-target-Cre line” combinations). Since thalamic areas are always the source in TC connections, the initial hierarchy score of each thalamic area *i* is defined by the average direction of connections from the area:

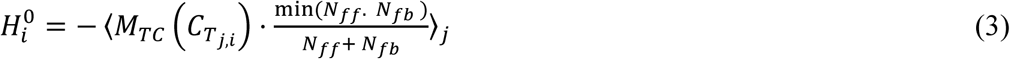

The parameters *N*_*ff*_ and *N*_*fb*_ refer to the numbers of feedforward and feedback thalamocortical connections, respectively. Once the initial positions of the thalamic areas in the hierarchy are obtained using Eq 3, hierarchy scores of thalamic and cortical areas are iterated until the fixed points are reached, using a full mapping function *M*_*CC*+*TC*_ that combines *M*_*CC*_ and *M*_*TC*_, as done with the cortical hierarchy based on CC connections only (Eq 2).

To test the significance of the hierarchy levels of these areas, we generated 100 sampled connectivity data of the same size via bootstrapping, and computed the hierarchy scores of the eight regions using the bootstrapped connectivity data. We performed Wilcoxon paired signed rank sum tests on these scores, showing that hierarchy levels of LM and RL cannot be meaningfully distinguished (*P* = 0.08) but the rest of the areas are at significantly distinct hierarchical positions, with the 5% confidence level.

#### 9.9. Other statistical methods

To quantify the correlation between the mean value of each metric and the anatomical hierarchy score, both the Pearson correlation coefficient (scipy.stats.pearsonr) and Spearman’s rank correlation coefficient (scipy.stats.spearmanr) were used.

To test for significant differences between pairs of areas, a Wilcoxon rank–sum statistic was used (scipy.stats.ranksum). For time to first spike, receptive field size, modulation index, and firing rate, each unit was considered an independent sample. For autocorrelation timescale, which is computed across all units in one area, each area for one mouse was considered an independent sample. Correction for multiple comparisons was performed using the Benjamini-Hochberg False Discovery Rate (statsmodels.stats.multitest.multipletests).

### 10. Data processing pipeline

Data for each session was uploaded to the Allen Institute Laboratory Information Management System (LIMS). Each dataset was run through the same series of processing steps using a set of project-specific workflows. Out of 61 sessions entering the processing pipeline, 58 resulted in successful NWB file generation. The 3 processing failures were due to mismatches in session identifiers or expected file structures that prevented the workflow from completing.

### 11. Data and code availability

The data from all 58 passive viewing experiments used to generate main text Figures 1 through 4 is available for download in Neurodata Without Borders format via the AllenSDK. Example Jupyter Notebooks for accessing the data can be found at https://allensdk.readthedocs.io/en/latest/visual_coding_neuropixels.html.

The metrics table used to generate Figure 5E–F and Supplementary Figure 10 is available in the GitHub repository for this manuscript (see below). The remaining data for the active behavior experiments will be made available upon request.

Code is available in the following repositories:

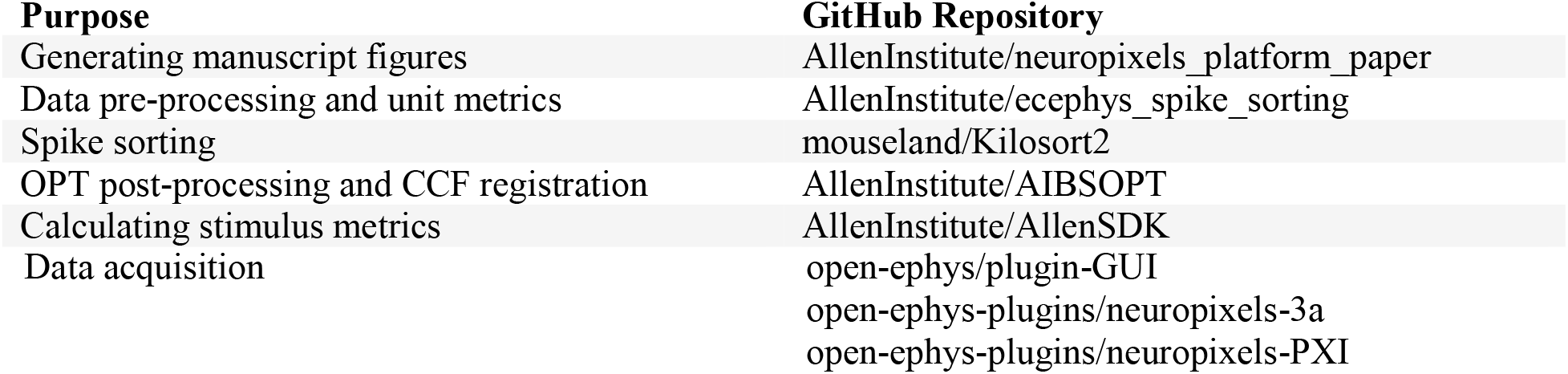

### 12. Open-source software libraries

NumPy (van der Walt et al., 2011)

SciPy (Jones et al., 2001)

IPython (Pérez and Granger, 2007)

Matplotlib (Hunter, 2007)

Pandas (McKinney, 2010)

xarray (Hoyer and Hamman, 2017)

scikit-learn (Pedregosa et al., 2012)

VTK (Schroeder et al., 2006)

DeepLabCut (Mathis et al., 2018; Nath et al., 2019)

statsmodels (Seabold and Perktold, 2010)

allenCCF (Shamash et al., 2018)

tifffile - https://pypi.org/project/tifffile/ Jupyter - https://jupyter.org/

pynwb - https://pynwb.readthedocs.io/en/stable/

